# Unmodified, autologous adipose-derived regenerative cells improve cardiac function, structure and revascularization in a porcine model of chronic myocardial infarction

**DOI:** 10.1101/286468

**Authors:** Alexander Haenel, Mohamad Ghosn, Tahereh Karimi, Jody Vykoukal, Claudia Kettlun, Dipan Shah, Amish Dave, Miguel Valderrabano, Daryl Schulz, Alon Azares, Albert Raizner, Eckhard Alt

## Abstract

Numerous studies have investigated cell-based therapies for myocardial infarction (MI), with mixed results. In the present study the left anterior descending (LAD) artery of pigs was occluded for 180 min. Four weeks later, the mean left ventricular ejection fraction (LVEF) was shown to have been reduced to approximately 35%. At that time, 18×106 unmodified, autologous adipose-derived regenerative cells (UA-ADRCs) were delivered into the LAD vein (control: delivery of saline). Six weeks following UA-ADRCs/saline delivery, the mean LVEF had increased by 18% (p<0.01) after delivery of UA-ADRCs, but was unchanged after delivery of saline. This is among the best outcome ever reported in studies on porcine animal models of cell-based therapies for MI in which functional outcome was assessed with cardiac magnetic resonance imaging. The unique combination of the procedure used for isolating UA-ADRCs, the late cell delivery time and the uncommon cell delivery route applied in the present study may open new horizons for cell-based therapies for MI.

## Introduction

Cardiovascular diseases substantially contribute to morbidity and mortality worldwide^1^. Myocardial infarction (MI) is one of the most common consequences of ischemic heart disease (IHD)^2^. Advanced medical treatments and device-based therapies have substantially improved the survival of patients with MI^3^. However, these therapies can only rescue the remaining viable myocardial tissue within the damaged heart, but not replace lost myocardium with regenerated cardiovascular cells. In this regard cell-based therapies have emerged as a promising strategy to regenerate ischemic myocardium^4-6^. However, clinical trials have shown substantial differences in efficacy and potency, and some trials failed to show any benefit^6^. Some authors speculated that the limited clinical success of cell-based therapies for MI is mainly due to ineffective homing of the cells to the heart and their lack of differentiation into cardiomyocytes^7^. The following explanations show, however, that the results of preclinical and clinical studies available so far justify the further optimization of cell-based therapies for MI, rather than giving up this promising form of therapy prematurely.

Intensive basic research over the last decade has shown that the following factors mainly influence the effects of cell-based therapies for MI: cell type, cell dose, cell delivery route, cell delivery timing and potential augmentation of cellular survival and/or function by physical stimulation, chemical/pharmacological treatment and genetic modification^6,8^. However, it is critical to note that particularly the selection of cell types and the augmentation of cellular survival and/or function are restricted by human safety issues^9-12^. For example, substantial safety concerns such as tumorigenesis severely limit the clinical translation of induced pluripotent stem cells^13,14^. Furthermore, clinical studies applying allogenic cells in cell-based therapies reported the production of donor-specific antibodies^15,16^, which is not the case when using autologous cells.

Cells derived from adipose tissue, either freshly isolated (named stromal vascular fraction (SVF) or adipose-derived regenerative cells (ADRCs)) or culture-expanded (named adipose-derived stem cells (ASCs)) have emerged as a promising tool for regenerating ischemic myocardium after MI^5,17^. Both ADRCs and ASCs have a number of advantages over other cell types in cell-based therapies for MI. For example, adipose tissue has a significantly higher stem cell density than bone marrow (5% vs. 0.01%), and harvesting adipose tissue is much less invasive and painful than harvesting bone marrow^5,18,19^. Unmodified ADRCs may be less efficacious than culture-expanded (and potentially modified) ASCs in regenerating ischemic myocardium, but have the advantage of lower safety requirements because no (*ex vivo*) culturing and no modification is involved, which also leads to substantially lower costs.

A number of enzymatic and non-enzymatic systems for isolating ADRCs were developed (for a non-up-to-date overview see^20^). The reported cell yield after some of these different procedures varied considerably^21^. Furthermore, it was shown that in general, enzymatic isolation of ADRCs yielded more cells than non-enzymatic (mechanical) isolation^22^.

With regard to cell delivery routes, a recent meta-analysis of preclinical studies and clinical trials on the efficacy of mesenchymal stem cell therapy for MI suggested superiority of transendocardial stem cell injection (TESI) over direct intramyocardial injection, intravenous infusion and intracoronary infusion^23^. However, TESI cannot be considered the optimum cell delivery route because the POSEIDON randomized clinical trial demonstrated that scar size reduction and ventricular functional responses preferentially occurred at the sites of TESI^24^.

Both ADRCs and ASCs were applied in a number of experimental studies using rodent, rabbit, porcine and sheep animal models^17^; details of studies using porcine and sheep animal models are summarized in Supplementary Table 1. Porcine and sheep models of experimentally induced transmural MI have the great advantage over rodent and rabbit models that potential therapeutic application of ADRCs and ASCs can be performed using the same instrumentation and standard of care as in humans^25^. Furthermore, porcine models allow the opportunity to evaluate the therapeutic outcome with MRI, which is considered to provide the most accurate, comprehensive and reproducible measurements of cardiac chamber dimensions, volumes, function and infarct size compared with other techniques^8^. Unfortunately, in those studies on porcine and sheep models in which treatment success of cell-based therapies for (experimentally induced) MI was assessed using MRI, temporary occlusion of the left anterior descending (LAD) artery was (except of one study that showed no success^26^) restricted to 60 min^27,28^ or 90 min^29,30^. As a result, in most of these studies^27,28,30^ the left ventricular ejection fraction (LVEF) after MI was between 42% and 54%, which may in clinical settings not represent the correct target population with moderate to severe left ventricular dysfunction^8^. Furthermore, cells were delivered immediately after experimental induction of MI in twelve out of the 16 preclinical studies on cell-based therapy for MI using porcine and sheep animal models summarized in Supplementary Table 1. However, studies on rats and pigs indicated that delivery of cells at a later time (i.e., after the acute inflammation period of MI) may yield more favorable outcome^31-33^.

Taking into account all these factors the aim of the present feasibility study was to determine (i) whether experimental occlusion of the LAD artery of pigs for three hours at time point T0 (Fig. 1a-c) results in a clinically relevant reduction of the LVEF to less than 40% at four weeks after T0 (i.e., at time point T1); and (ii) whether the delivery of a clinically relevant dose of unmodified, autologous (UA) ADRCs (18 × 10^6^ cells) into the temporarily blocked LAD vein (Fig. 1d) at T1 is a safe treatment that results in MRI-demonstrated improved cardiac function and structure ten weeks after T0 (i.e., six weeks after T1) compared to the situation at T1.

**Fig. 1.**
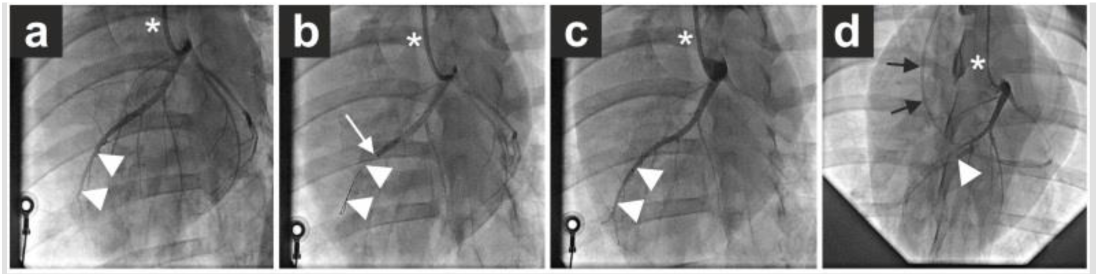
Experimental details of the present study. **a** Baseline coronary angiogram of a porcine heart in a left anterior oblique view (in all panels the asterisk indicates the angiography catheter positioned in the left main coronary ostium). The white arrowheads indicate the distal left anterior descending (LAD) artery. **b** Induction of myocardial infarction by occlusion of the LAD artery for three hours through an inflated balloon catheter at time point T0. The white arrow indicates the position of the inflated balloon inside the mid LAD artery, whereas the white arrowheads show the guidewire in the distally occluded LAD artery. **c** Complete reperfusion of the LAD artery (white arrowheads) three hours after removal of the balloon occlusion. **d** Delivery of unmodified, autologous adipose-derived regenerative cells (or saline as control, respectively) through the LAD vein (matching the initial LAD artery occlusion site) into the infarction area four weeks later (i.e., at time point T1). To this end, the LAD vein was occluded with another inflated balloon catheter that was advanced through a guiding catheter (black arrows), placed from the right jugular vein into the right atrium and then into the coronary sinus. The inflated balloon (filled with contrast dye; white arrowhead) in the coronary vein had the aim to prevent backflow of cells when they were delivered through the distal orifice of the central lumen of this balloon catheter. When more cells were injected, the pressure in the compartment between the blocked distal capillary passage and the proximal inflated balloon in the coronary vein was slightly increased.

To test these hypotheses, UA-ADRCs were isolated from adipose tissue that was extracted from the nuchal region of the pigs, and were characterized with flow cytometry. Processing of extracted adipose tissue to the ready-to-inject UA-ADRCs suspension was performed with the Matrase enzyme blend and the Transpose RT system (both from InGeneron, Inc., Houston, TX, USA). Delivery of saline was used as control. Effects of UA-ADRCs were assessed with electrocardiogram-gated, steady-state free precession cardiac magnetic resonance (SSFP CMR) imaging (Fig. 2), late gadolinium enhancement (LGE) CMR imaging and detailed post mortem histologic analysis. A comprehensive timeline of the present study is provided in Table 1.

**Table 1.**
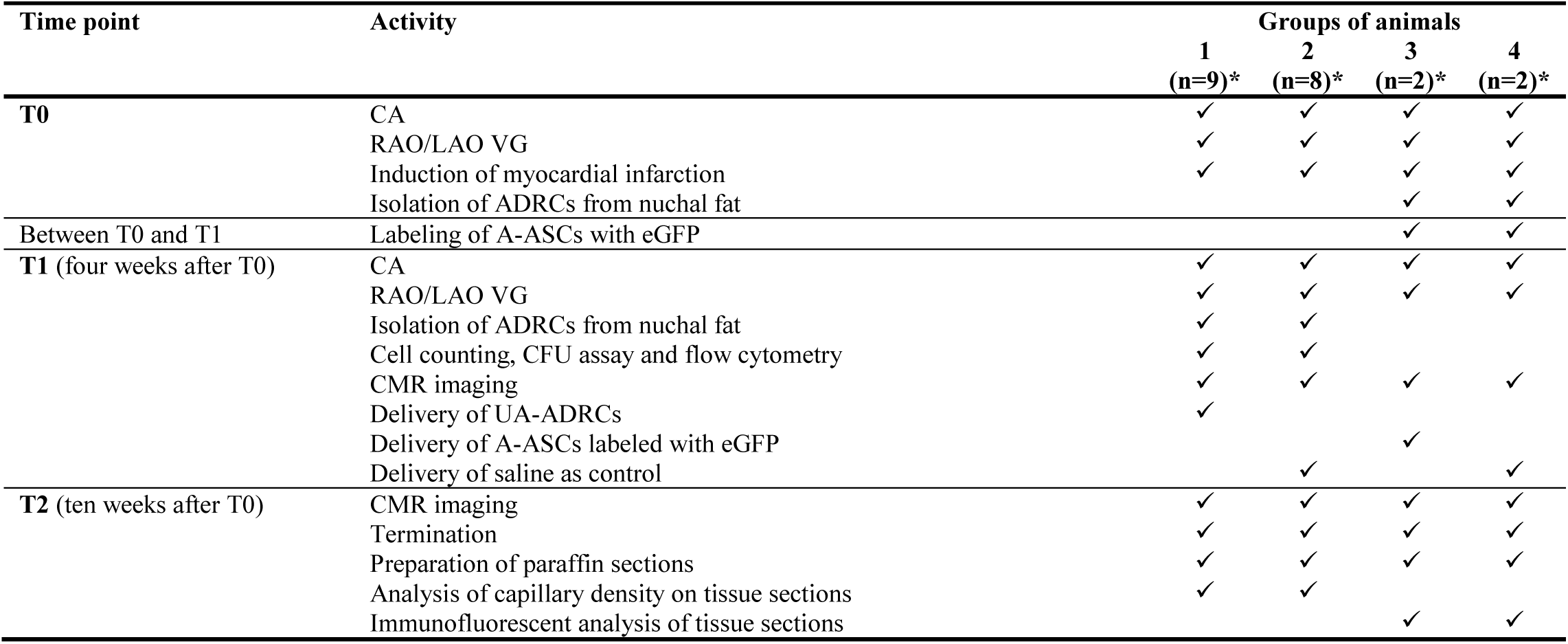
Timeline of the present study. Details are provided in the main text. Abbreviations: CA, coronary angiography; RAO/LAO VG, ventriculography in right and left anterior oblique views; ADRCs, adipose-derived regenerative cells; A-ASCs, autologous adipose-derived stem cells; CFU assay, colony forming unit assay; UA-ADRCs, unmodified, autologous adipose-derived regenerative cells; CMR, cardiac magnetic resonance; *, numbers of animals in each group that were included in the final analysis (details are provided in the Methods section).

| Time point | Activity | Groups of animals |  |  |  |
| --- | --- | --- | --- | --- | --- |
|  |  | 1<br>(n=9)* | 2<br>(n=8)* | 3<br>(n=2)* | 4<br>(n=2)* |
| **T0** | CA | ✓ | ✓ | ✓ | ✓ |
|  | RAO/LAO VG | ✓ | ✓ | ✓ | ✓ |
|  | Induction of myocardial infarction | ✓ | ✓ | ✓ | ✓ |
|  | Isolation of ADRCs from nuchal fat |  |  | ✓ | ✓ |
| Between T0 and T1 | Labeling of A-ASCs with eGFP |  |  | ✓ | ✓ |
| **T1** (four weeks after T0) | CA | ✓ | ✓ | ✓ | ✓ |
|  | RAO/LAO VG | ✓ | ✓ | ✓ | ✓ |
|  | Isolation of ADRCs from nuchal fat | ✓ | ✓ |  |  |
|  | Cell counting, CFU assay and flow cytometry | ✓ | ✓ |  |  |
|  | CMR imaging | ✓ | ✓ | ✓ | ✓ |
|  | Delivery of UA-ADRCs | ✓ |  |  |  |
|  | Delivery of A-ASCs labeled with eGFP |  |  | ✓ |  |
|  | Delivery of saline as control |  | ✓ |  | ✓ |
| **T2** (ten weeks after T0) | CMR imaging | ✓ | ✓ | ✓ | ✓ |
|  | Termination | ✓ | ✓ | ✓ | ✓ |
|  | Preparation of paraffin sections | ✓ | ✓ | ✓ | ✓ |
|  | Analysis of capillary density on tissue sections | ✓ | ✓ |  |  |
|  | Immunofluorescent analysis of tissue sections |  |  | ✓ | ✓ |

**Fig. 2.**
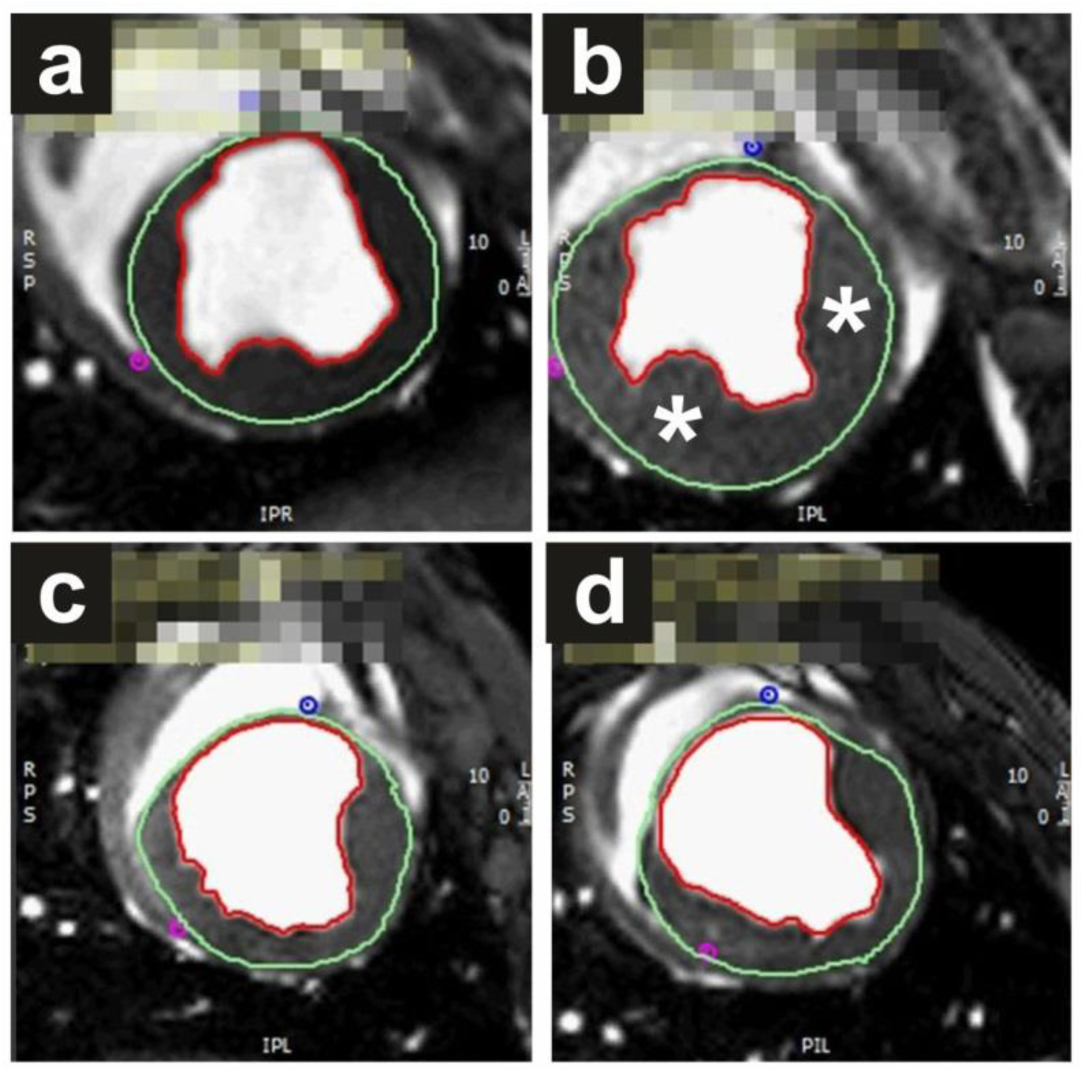
Representative examples of end-systolic, short axis, transversal images through the mid left ventricle of a porcine heart obtained with steady-state free precession cardiac magnetic resonance (SSFP CMR) imaging for analyzing hemodynamic parameters and wall motility at four weeks (time point T1) (**a**,**c**) and ten weeks (time point T2) (**b**,**d**) after experimental occlusion of the left anterior descending (LAD) artery by means of a balloon catheter for three hours at time point T0, followed by the delivery of respectively unmodified, autologous adipose-derived regenerative cells (UA-ADRCs) (**a**,**b**) or saline as control (**c**,**d**) into the LAD vein (matching the initial LAD occlusion site) immediately after SSFP CMR imaging at T1. In all panels the epicardial contours are highlighted in green, and the endocardial contours in red. Note the increased end-systolic thickness of the left ventricular wall at T2 after delivery of UA-ADRCs at T1 (asterisks in **b**) compared to the delivery of saline at T1 (**d**). In the examples shown here the left ventricular ejection fraction was 27.2% in **a**, 39.7% in **b**, 22.5% in **c** and 27.2% in **d**. Corresponding cine sequences are provided in Supplementary Movies 1 to 4.

Because no preliminary data were available, the null hypothesis of the present study was that the delivery of UA-ADRCs into the temporarily blocked LAD vein four weeks after experimentally induced MI is not an effective treatment for MI in a porcine animal model.

## Results

### Unmodified, autologous adipose-derived regenerative cells expressed markers indicating potential for tissue regeneration

On average 18.1 ± 1.61 g (mean ± standard error of the mean; SEM) (range, 12-25) of adipose tissue were obtained from the nuchal region of each pig. The average yield of nucleated cells obtained after processing was 0.98 ± 0.10 x 10^6^ cells per gram adipose tissue. Cell viability before delivery (determined with the Trypan Blue exclusion assay) was 93.3 ± 0.4%. Summarized results of flow cytometric analysis (i.e., mean relative number of cells immunopositive for a certain marker) of the fresh, uncultured UA-ADRCs were as follows: CD29: 48.5% (range, 44.1-52.9); CD44: 37.0% (34.2-39.8); CD31: 11.4% (6.0-16.7); NG2: 9.78% (7.25-12.3); CD45: 8.92% (3.9-13.9); Oct4: 4.3% (2.4-6.2); Nestin: 4.1% (1.9-6.2); CD146: 0.6% (0.4-0.7); and CD117: 0.3% (0.1-0.4) (an example of the results of flow cytometric analysis is shown in Suppl. Fig. 1). Colony formation as an indicator of stemness was 11.3% (assessed with a colony forming unit [CFU] assay).

### Improved cardiac function after delivery of unmodified, autologous adipose-derived regenerative cells

Unlike the animals in Group 2 (delivery of saline), the animals in Group 1 (delivery of UA-ADRCs) showed the following, statistically significant improvements in cardiac function at T2 compared to T1: increased mean cardiac output (+38%) (Fig. 3a,b), increased mean stroke volume (+41%) (Fig. 3c,d), and increased mean left ventricular ejection fraction (+18%) (Fig. 3e,f) (mean and SEM data are also provided in Supplementary Table 2). Accordingly, the null hypothesis was rejected.

**Fig. 3.**
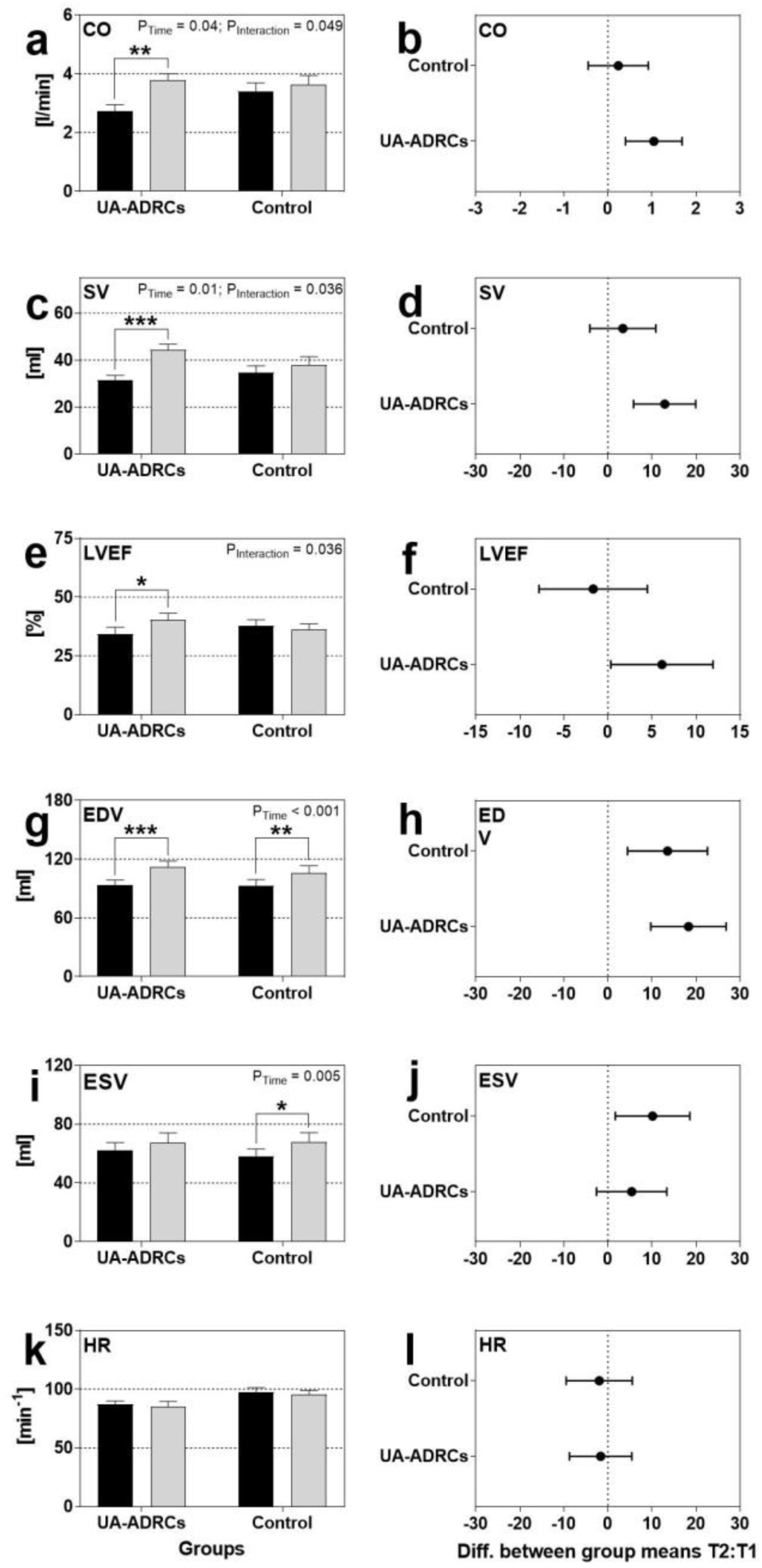
Functional alterations of the porcine heart (investigated with steady-state free precession cardiac magnetic resonance imaging) at ten weeks (i.e., at time point T2) after experimental occlusion of the left anterior descending (LAD) artery for three hours at time point T0, followed by the delivery of respectively 18 × 10^6^ unmodified, autologous adipose-derived regenerative cells (UA-ADRCs) (n=9) or saline (Control) (n=8) into the balloon-blocked LAD vein (matching the initial LAD artery occlusion site) at four weeks after T0 (i.e., at time point T1). **a**,**c**,**e**,**g**,**i**,**k** Group-specific mean and standard error of the mean (SEM) of cardiac output (CO; **a**), stroke volume (SV; **c**), left ventricular ejection fraction (LVEF; **e**), end-diastolic volume (EDV; **g**), end-systolic volume (ESV; **i**) and heart rate (HR; **k**) immediately before T1 (closed bars) and at T2 (gray bars). Except of the end-systolic volume at T2 of the animals treated with UA-ADRCs all data passed the Shapiro-Wilk normality test. P-values of repeated measures two-way analysis of variance smaller than 0.05 are provided in the upper right corner of the panels. Results of group-specific Bonferroni’s multiple comparison tests are indicated (*, p<0.05; **, p<0.01; ***, p<0.001). **b**,**d**,**f**,**h**,**j**,**l** 95% confidence intervals (Bonferroni) of the differences of group-specific mean data between T2 and T1.

The mean end-diastolic volume was statistically significantly increased in both groups at T2 compared to T1 (Group 1: +20%; Group 2: +15%) (Fig. 3g,h), whereas the mean end-systolic volume was statistically significantly increased only in Group 2 (+17%) but not in Group 1 (Fig. 3i,j). The mean heart rate showed no statistically significant difference between T1 and T2 (Fig. 3k,l).

### Improved cardiac structure after delivery of unmodified, autologous adipose-derived regenerative cells

Unlike the animals in Group 2, the animals in Group 1 showed a statistically significantly increased mean mass of the left ventricle at T2 compared to T1 (+29%) (Fig. 4a,b). Besides this, the mean relative amount of scar volume of the left ventricular wall was statistically significantly increased in Group 2 at T2 compared to T1 (+29%), but was statistically significantly decreased in Group 1 at T2 compared to T1 (-21%) (Fig. 4c,d and Fig. 5). Analysis of regional wall thickening and regional replacement fibrosis is shown in Suppl. Fig. 2.

**Fig. 4.**
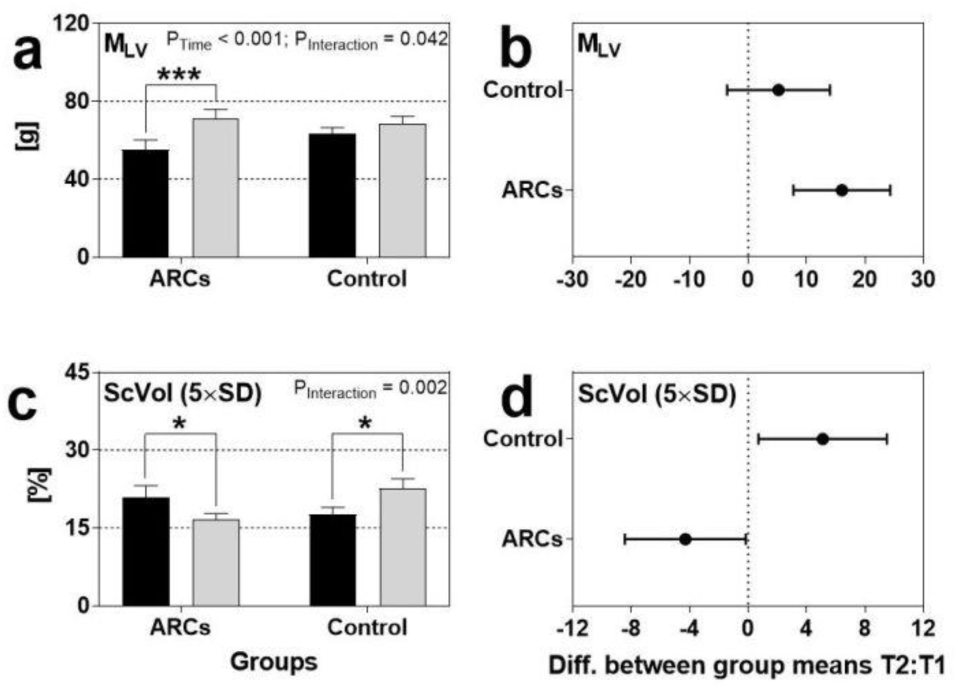
Structural alterations of the porcine heart at ten weeks (i.e., at time point T2) after experimental occlusion of the left anterior descending (LAD) artery for three hours at time point T0, followed by the delivery of respectively 18 × 10^6^ unmodified, autologous adipose-derived regenerative cells (UA-ADRCs) (n=9) or saline (Control) (n=8) into the balloon-blocked LAD vein (matching the initial LAD artery occlusion site) at four weeks after T0 (i.e., at time point T1). **a**,**c** Group-specific mean and standard error of the mean (SEM) of the left ventricular mass (investigated with steady-state free precession cardiac magnetic resonance [CMR] imaging) (M_LV_; **a**) and the relative amount of scar volume of the left ventricular wall (investigated with late gadolinium enhancement CMR imaging) (ScVol; **c**) immediately before T1 (closed bars) and at T2 (gray bars). All data passed the Shapiro-Wilk normality test. P values of repeated measures two-way analysis of variance smaller than 0.05 are provided in the upper right corner of the panels. Results of group-specific Bonferroni’s multiple comparison tests are indicated (*, p<0.05; ***, p<0.001). **b**,**d** 95% confidence intervals (Bonferroni) of the differences of group-specific mean data between T2 and T1.

**Fig. 5.**
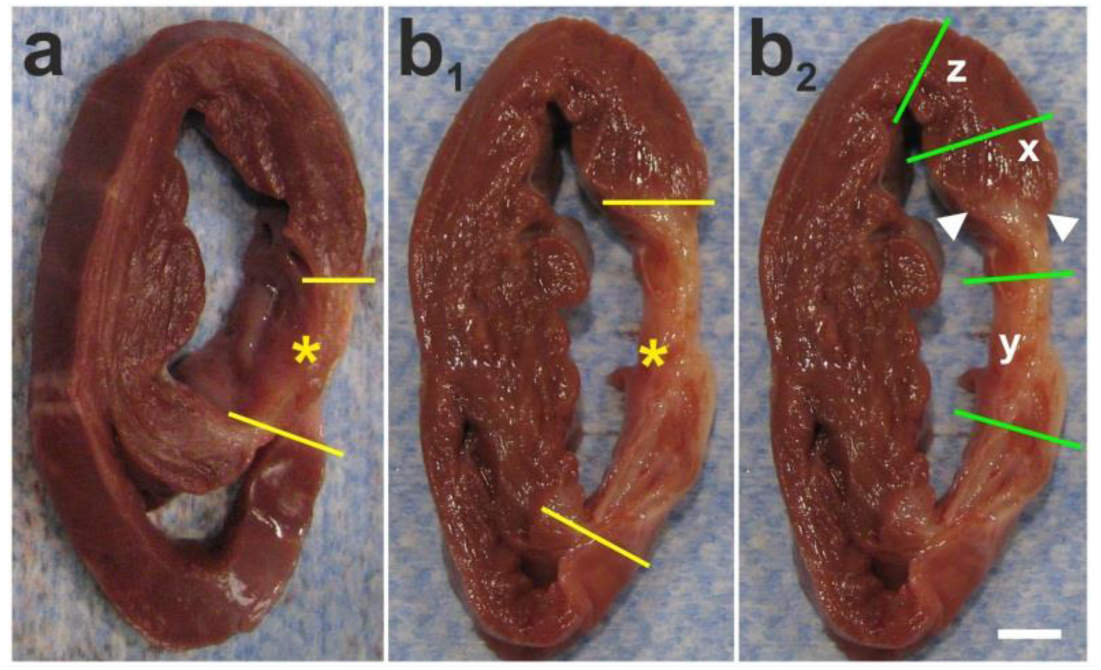
Representative, transversal, 1 cm-thick slices of post mortem porcine hearts at ten weeks (i.e., at time point T2) after experimental occlusion of the left anterior descending (LAD) artery for three hours at time point T0, followed by the delivery of respectively 18 × 10^6^ unmodified, autologous adipose-derived regenerative cells (**a**) or saline (**b**) into the balloon-blocked LAD vein (matching the initial LAD artery occlusion site) at four weeks after T0 (i.e., at time point T1). The yellow lines in **a** and **b**_**1**_ indicate the left ventricular border zones of the myocardial infarction (MI) (yellow asterisks in **a**,**b**_**1**_). The relative amount of scar volume of the left ventricular wall was 15.4% at T1 and 11.9% at T2 in **a**, and 10.2% at T1 and 19.1% at T2 in **b. b**_**2**_ same panel as in b_1_, indicating the generation of tissue blocks containing the left ventricular border zone of MI (x; the borderzone is indicated by white arrowheads), the core region of MI (y) and regions of viable myocardium (z). The scale bar in **b**_**2**_ represents 1 cm in **a**,**b**.

At the microscopic level the left ventricular border zone of the myocardial infarction of the post mortem hearts from the animals in Group 1 showed patchy islets of cardiomyocytes located within areas of fibrous tissue, whereas the same region in the post mortem hearts from some animals in Group 2 showed infiltration with inflammatory cells (Fig. 6).

**Fig. 6.**
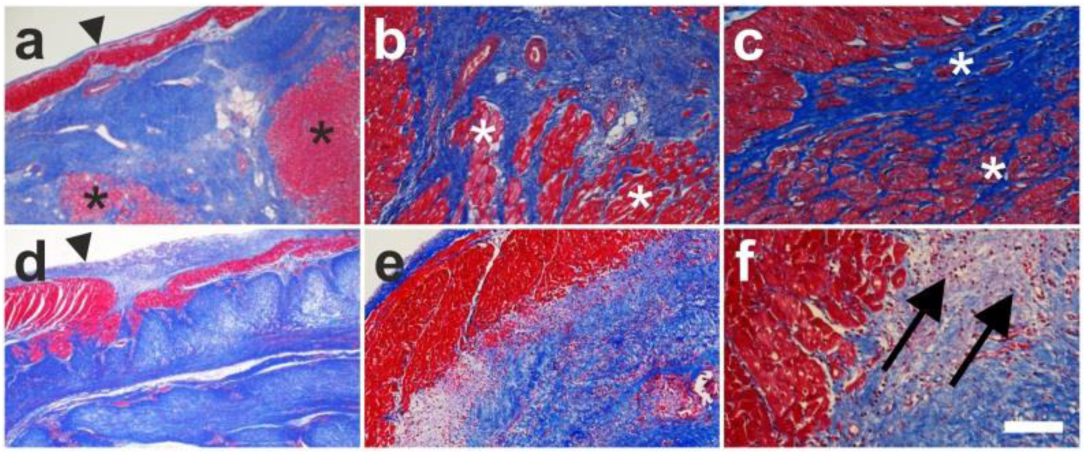
Representative photomicrographs of paraffin-embedded, 5 µm thick tissue sections of post mortem hearts from pigs, taken from the left ventricular border zone of myocardial infarction at ten weeks (i.e., at time point T2) after experimental occlusion of the left anterior descending (LAD) artery for three hours at time point T0, followed by the delivery of respectively 18 × 10^6^ unmodified, autologous adipose-derived regenerative cells (**a**-**c**) or saline as control (**d**-**f**) into the balloon-blocked left LAD vein (matching the initial LAD artery occlusion site) at four weeks after T0 (i.e., at time point T1). The sections were stained with Masson’s Trichrome staining. Collagen is stained blue and cardiomyocytes are stained red. The arrowheads in **a**,**d** point to the endocard, the asterisks in **a**-**c** indicate patchy islets of cardiomyocytes located within areas of fibrous tissue, and the arrows in **f** point to an infiltration with inflammatory cells. The scale bar in **f** represents 500 µm in **a**,**d**, 200 µm in **b**,**e**, and 100 µm in **c**,**f**.

### Improved cardiac revascularization after delivery of unmodified, autologous adipose-derived regenerative cells

The left ventricular border zone of the myocardial infarction of the post mortem hearts from the animals in Group 1 showed a statistically significantly higher mean microvessel density than corresponding sections of the post mortem hearts from the animals in Group 2 at T2 (54.4 ± 3.7 vs. 26.1 ± 2.8 [mean ± SEM] capillaries per mm^2^; p < 0.001) (Fig. 7a,c). Immunofluorescent detection of von Willebrand factor supported this finding (Fig. 7b,d).

**Fig. 7.**
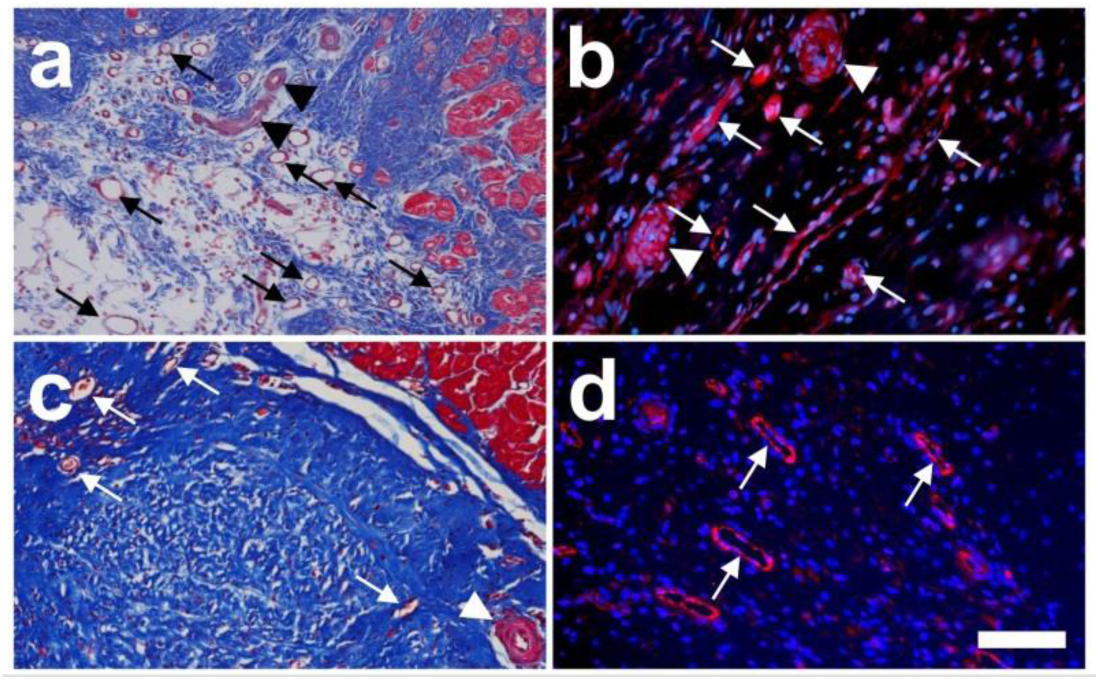
Representative photomicrographs of paraffin-embedded, 5 µm thick tissue sections of post mortem hearts from pigs, taken from the left ventricular border zone of myocardial infarction at ten weeks (i.e., at time point T2) after experimental occlusion of the left anterior descending (LAD) artery for three hours at time point T0, followed by the delivery of respectively 18 × 10^6^ unmodified, autologous adipose-derived regenerative cells (**a**,**b**) or saline as control (**c**,**d**) into the balloon-blocked LAD vein (matching the initial LAD artery occlusion site) at four weeks after T0 (i.e., at time point T1). The sections were stained with Masson’s Trichrome staining (**a**,**c**) or processed for immunofluorescent detection of von Willebrand factor (red) and counterstained with DAPI (blue) (**b**,**d**). In (**a**,**c**) collagen is stained blue and cardiomyocytes are stained red. The arrows point to microvessels and the arrowheads to small arterioles. The scale bar in **d** represents 100 µm in **a**,**c** and 35 µm in **b**,**d**.

### Proliferation of autologous adipose-derived stem cells in vivo and integration into the wall of microvessels in the left ventricular border zone of the myocardial infarction

Autologous adipose-derived stem cells (A-ASCs) that were (i) isolated from porcine nuchal fat at the time point of experimental occlusion of the LAD artery, (ii) labeled *in vitro* with enhanced green fluorescent protein (eGFP) and (iii) delivered into the LAD vein four weeks later (animals in Group 3) were found in the wall of microvessels in the left ventricular border zone of the myocardial infarction at T2 (Figs. 8 and 9). Some of these cells were immunopositive for Ki-67, indicating that they had re-entered the cell cycle (Fig. 10).

**Fig. 8.**
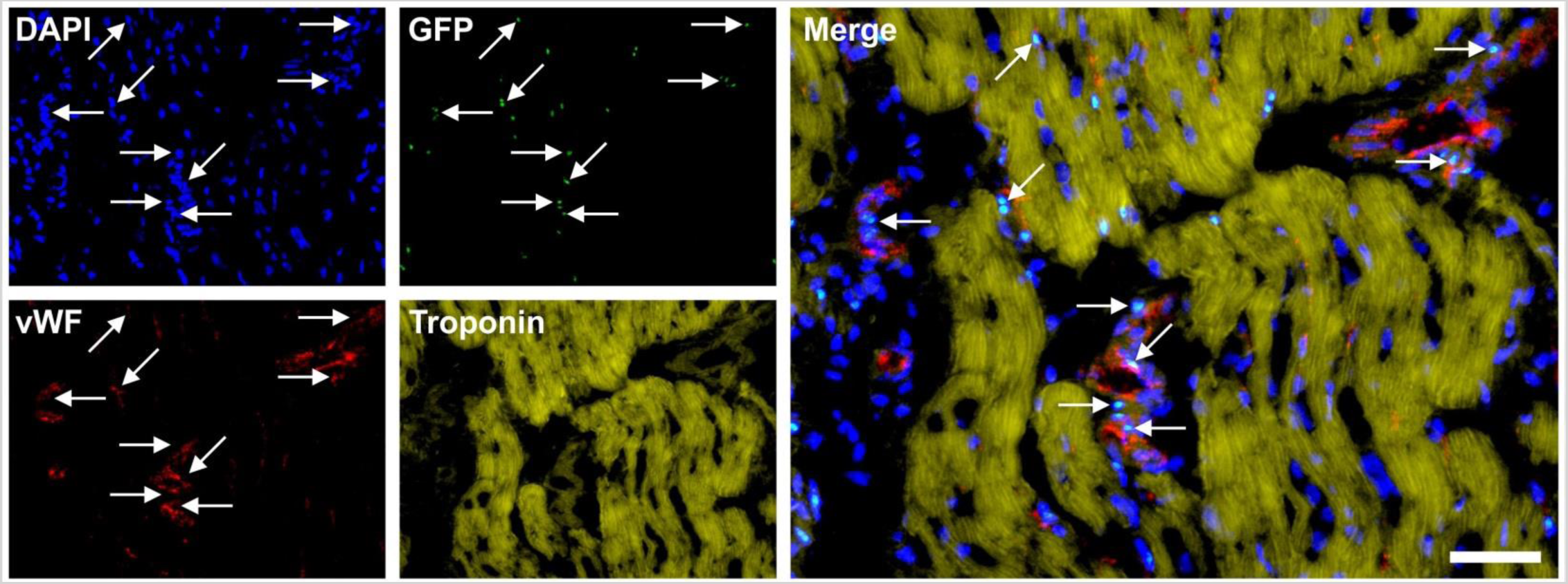
Representative photomicrographs of a paraffin-embedded, 5 µm thick tissue section of a post mortem heart from a pig, taken from the left ventricular border zone of myocardial infarction ten weeks after experimental occlusion of the left anterior descending (LAD) artery for three hours, followed by the delivery of eGFP-labeled autologous adipose-derived stem cells into the balloon-blocked LAD vein (matching the initial LAD occlusion site) at four weeks after occlusion of the LAD. The section was stained with DAPI (blue) and processed for immunofluorescent detection of GFP (green), von Willebrand factor (vWF) (red) and Troponin (yellow). The arrows indicate cell nuclei that were immunopositive for GFP and were found in the wall of small vessels (the positions of these cell nuclei are also labeled in the panel representing vWF). The scale bar represents 50 µm in the merged panel, and 100 µm in the individual panels.

**Fig. 9.**
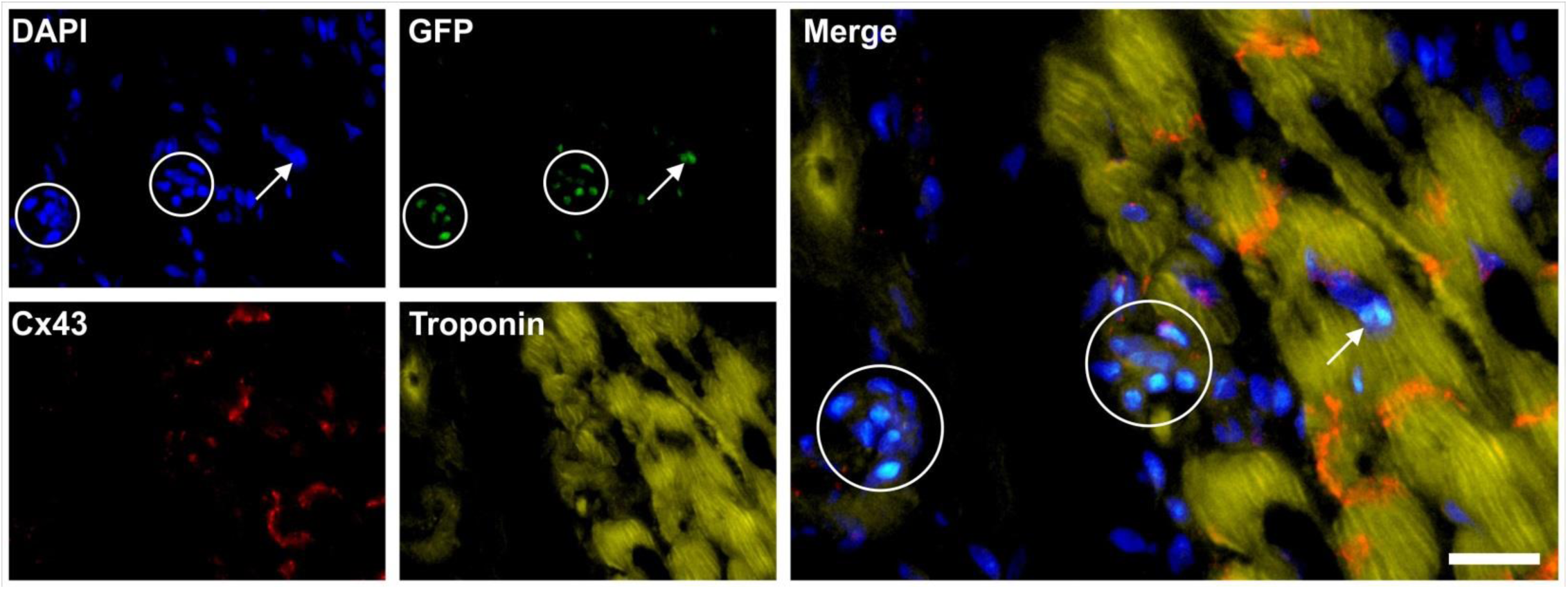
Representative photomicrographs of a paraffin-embedded, 5 µm thick tissue section of a post mortem heart from a pig, taken from the left ventricular border zone of myocardial infarction ten weeks after experimental occlusion of the left anterior descending (LAD) artery for three hours, followed by the delivery of eGFP-labeled autologous adipose-derived stem cells into the balloon-blocked LAD vein (matching the initial LAD occlusion site) at four weeks after occlusion of the LAD. The section was stained with DAPI (blue) and processed for immunofluorescent detection of GFP (green), Cx43 (red) and Troponin (yellow). The circles indicate regions where most of the cell nuclei were immunopositive for GFP, and the arrow a GFP-positive cell nucleus inside a cardiomyocyte. The scale bar represents 25 µm in the merged panel, and 50 µm in the individual panels.

**Fig. 10.**
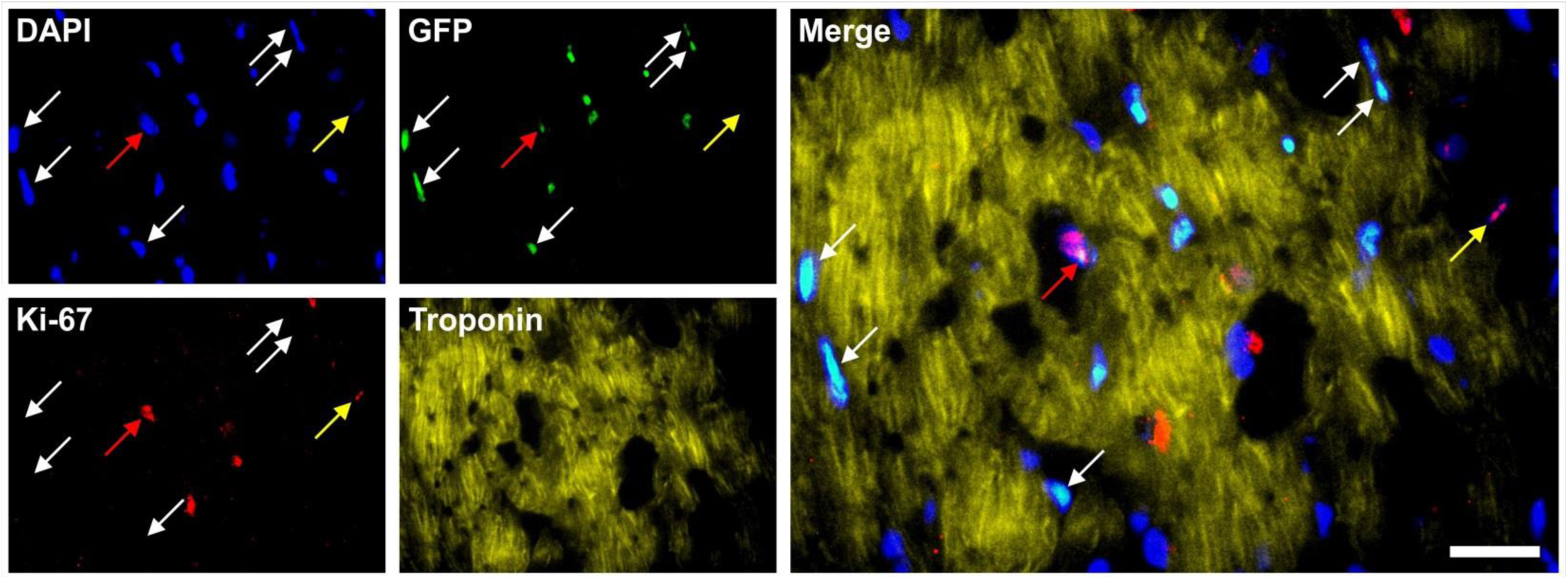
Representative photomicrographs of a paraffin-embedded, 5 µm thick tissue section of a post mortem heart from a pig, taken from the left ventricular border zone of myocardial infarction ten weeks after experimental occlusion of the left anterior descending (LAD) artery for three hours, followed by the delivery of eGFP-labeled autologous adipose-derived stem cells into the balloon-blocked LAD vein (matching the initial LAD occlusion site) at four weeks after occlusion of the LAD. The section was stained with DAPI (blue) and processed for immunofluorescent detection of GFP (green), Ki-67 (red) and Troponin (yellow). The white arrows point to cell nuclei that were immunopositive for GFP but not for Ki-67, the yellow arrows to a cell nucleus that was immunopositive for Ki-67 but not for GFP, and the red arrows to a cell nucleus that was immunopositive for both GFP and Ki-67. The scale bar represents 25 µm in the merged panel, and 50 µm in the individual panels.

### No evidence of differentiation of unmodified, autologous adipose-derived regenerative cells into adipocytes

Cells in the left ventricular border zone of the myocardial infarction of the post mortem hearts from the animals in Groups 1 and 2 showed no expression of adiponectin at T2 (Fig. 11).

**Fig. 11.**
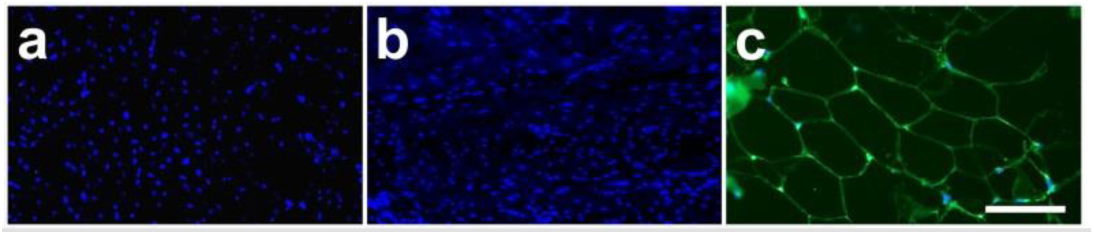
**a,b** Representative photomicrographs of paraffin-embedded, 5 µm thick tissue sections of post mortem hearts from pigs, taken from the left ventricular border zone of myocardial infarction ten weeks after experimental occlusion of the left anterior descending (LAD) artery for three hours, followed by the delivery of autologous, adipose-derived regenerative cells (**a**) or saline as control (**b**) into the balloon-blocked LAD vein (matching the initial LAD occlusion site) at four weeks after occlusion of the LAD. **c** Representative photomicrograph of a paraffin-embedded, 5 µm thick tissue section of subcutaneous adipose tissue from a pig. The sections were stained with DAPI (blue) and processed for immunofluorescent detection of adiponectin (green). The scale bar represents 100 µm.

## Discussion

During the last two decades, cell-based therapies for MI have been investigated in numerous pre-clinical studies and clinical trials, using (i) different types of cells (including ADRCs, ASCs, embryonic stem cells, induced pluripotent stem cells, bone marrow mononuclear cells, mesenchymal stem cells and cardiac progenitor cells), (ii) different animal models (including mouse, rat, pig, sheep and nonhuman primates) and during first clinical trials on patients, (iii) different pretreatments of cells (including physical stimulation, chemical/pharmacological treatment and genetic modification), (iv) different cell doses, (v) different cell delivery timing (including delivery of cells immediately after MI or at a later time after the acute inflammation period of MI), (vi) different cell delivery routes (including intracoronary and intravenous infusion, intramyocardial and transendocardial injection), and (vii) different patient inclusion and follow-up criteria. From the most recent reviews of these studies, the following, partially overlapping trends can be gleaned: (i) That enthusiasm and expectations should be dampened in connection with the potential for cell-based therapies for MI^34^; (ii) That the focus for further research should be on the optimal cell type, dose, delivery route and timing, as well as on precise mechanisms of action and long-term cell engraftment^6^; and (iii) That further research on potential new therapeutic strategies for MI should be considered, including *ex vivo* cell-mediated gene therapy, the use of biomaterials and cell-free therapies as well as stimulation of endogenous mechanisms of regeneration^7,8^.

The present study is the first that combines all of the following clinically relevant findings from research into cell-based therapies for MI: (i) the use of a porcine animal model that allows the study of cell-based therapies for chronic MI using the same instrumentation and standard of care as in humans (rather than using mice or rats); (ii) the experimental occlusion of the LAD artery for three hours (rather than for only 45 to 90 min); (iii) the requirement of a baseline LVEF after induction of MI of approximately 35% (rather than a clinically less relevant baseline LVEF of approximately 50%); (iv) the use of UA-ADRCs (rather than using ASCs, bone marrow-derived stem cells [BMSCs] or modified ASCs/BMSCs, respectively); (v) the delivery of UA-ADRCs at four weeks after induction of MI (rather than cell delivery immediately after MI); (vi) the delivery of UA-ADRCs retrograde into the temporarily blocked LAD vein (rather than transendocardial, intramyocardial or intracoronary cell delivery); and (vii) the demonstration of therapeutic success using CMR imaging (rather than using other methods including single photon emission computed tomography). Accordingly, the results of the present study may have the most direct relevance to the clinical practice.

Perhaps the most relevant findings of the present study ar the following: the statistically significant increase of the mean LVEF (+18% in relative terms) after delivery of UA-ADRCs was among the highest ever reported in studies on porcine animal models of cell-based therapies for MI in which functional outcome was assessed with CMR imaging (Fig. 12; details are provided in Supplementary Table 1). This was accompanied by a statistically significant decrease (-21%) in the mean relative amount of scar volume of the left ventricular wall (mScVol) at six weeks after delivery of UA-ADRCs (i.e., ten weeks after LAD occlusion), compared to +29% after delivery of saline. In this regard it is of note that the criterion applied for quantification of scar tissue (defined as hyper-enhancement of signal intensity in late gadolinium enhancement CMR imaging ≥ 5 standard deviations greater than normal myocardium^35^) was the strictest one among all studies addressed in Fig. 12 (details are provided in Supplementary Table 1). Accordingly, the mScVol data reported in the present study are not directly comparable to data reported in the other studies addressed in Fig. 12.

**Fig. 12.**
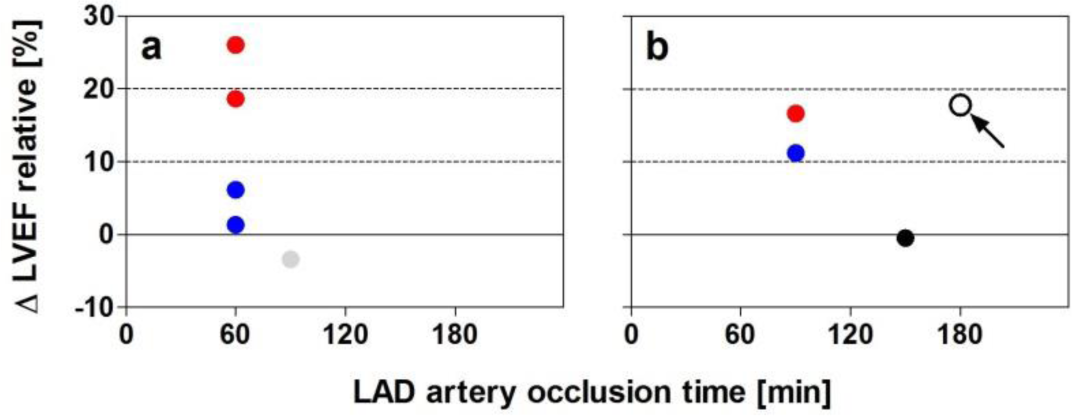
Change of the left ventricular ejection fraction (in relative terms; ΔLVEF) as a function of the occlusion time of the left anterior descending artery in those porcine animal models on cell-based therapies for myocardial infarction (MI) in which functional outcome was assessed with MRI. **a** Studies with delivery of cells immediately after induction of MI. **b** Studies with delivery of cells at a later time (between one and four weeks after MI). Cell-based therapy was performed with unmodified, autologous adipose-derived regenerative cells (open dot) in the present study (arrow); adipose-derived stem cells (ASCs) (blue dots in **a**,**b**), modified ASCs (red dots in **a**,**b**), bone marrow-derived stem cells (gray dot in **a**) and cardiosphere-derived stem cells (black dot in **b**). Details of these studies are summarized in Supplementary Table 1.

These results are particularly relevant because in the present study, the LAD occlusion time was the longest and the baseline LVEF at time of treatment was the lowest among the studies addressed in Fig. 12. Besides this, the present study was the only one among the studies addressed in Fig. 12 in which the cells were neither selected (as in case of ASCs) nor physically, chemically and/or genetically modified. We hypothesize that this was based on the unique combination of the enzymatic isolation procedure of cells using Matrase, the late cell delivery time and the novel cell delivery route applied in the present study.

The outcome of any cell-based therapy for MI critically depends on the delivered type of cells, which includes the isolation procedure in case of ADRCs. In the present study, cells were enzymatically isolated from adipose tissue using Matrase and the Transpose RT system (both from InGeneron). Matrase is a proprietary, optimized enzyme blend of collagenase I, collagenase II and a recombinantly generated neutral protease produced under GMP conditions. The Transpose RT system provides processing of adipose tissue together with Matrase in a predefined processing, washing and concentration cycle at 39° C within one hour.

The cells delivered in the present study were characterized as follows: (i) on average 49% of the cells were immunopositive for CD29 (CD29^+^) and on average 37% were CD44^+^. Both CD29 and CD44 are markers of ADRCs^17,21^. A similar result for CD29 was reported by Mitchell et al.^36^ when investigating human ADRCs (on average 48% of CD29^+^ cells), whereas results for CD44 were higher in Mitchell et al.^36^ (on average 64% of CD29^+^ cells) than in the present study. This may indicate species-specific differences in the expression of CD44. Mitchell et al.^36^ also reported that from passages 1 through 4, the mean relative number of CD29^+^ cells increased from 71% to 95%, and the mean relative number of CD44^+^ cells from 84% to 98% ^36^. (ii) On average only 9% of the cells were CD45^+^, a marker of blood derived cells^21,37^. This was substantially less than what was reported by Alt et al.^38^ in an earlier study on porcine ADRCs (on average 31%) as well as for human ADRCs (approximately 30%)^36,39^, and may reflect differences in the isolation procedure of the cells (Matrase was not used in^36,38,39^). (iii) On average 11% of the cells were CD31^+^, a marker of endothelial cells^21,39^. This was similar to earlier reports of porcine ADRCs (on average 8% in ^38^), whereas in studies on human ADRCs the mean relative number of CD31^+^ cells was reported as 1-6%^39^ or 22%^36^. The latter discrepancy may again reflect differences in the isolation procedure of ADRCs. (iv) The transcription factor Oct4 is highly associated with the pluripotency of stem cells but was also found expressed in human ASCs at passage 1^40^. In the present study, on average 4% of the delivered cells were Oct4^+^. On the other hand, Oct4 was not investigated in the other studies discussed here^17,21,36-39^, and relative numbers of Oct4^+^ ASCs at passage 1 were not reported^40^. (v) NG2 and CD146 are markers of pericytes^21,41^, Nestin is an early marker of neural stem/progenitor cells as well as of proliferative endothelial cells^42^, and CD117 (c-Kit) was described as a marker of hematopoietic progenitor cells, multipotent progenitors and common myeloid progenitors^43^. The mean relative numbers of cells delivered in the present study that were NG2^+^, CD146^+^, Nestin^+^ and CD117^+^ were 10%, 1%, 4% and 0.3%, respectively. Unfortunately, Alt et al.^38^ did not investigate the relative amount of NG2^+^, CD146^+^, Nestin^+^, and CD117^+^ cells in porcine ADRCs, and Mitchell et al.^36^ only reported the relative number of CD146^+^ cells in human ADRCs (on average 21%). Accordingly, one cannot determine whether the difference in the relative number of CD146^+^ cells reported in the present study and in the study by Mitchell et al.^36^ reflects species-specific differences, differences in the isolation procedure of ADRCs, or a mixture thereof.

Collectively, these data clearly indicate that ADRCs differ from ASCs and, thus, the outcome of studies in which ADRCs were delivered might not be directly compared to the outcome of studies in which ASCs were delivered. Furthermore, ADRCs used in different studies may show species-specific differences and in particular substantial differences caused by the isolation procedure of the ADRCs, even between different enzymatic isolation procedures (c.f. also^20-22^). In other words, the term ‘treatment with ADRCs’ can only be used as an umbrella term.

No other study on animal models assessed the therapeutic outcome of delivery of ADRCs in experimentally induced MI with CMR imaging. Furthermore, ADRCs were only tested in three clinical trials on cell-based therapy for cardiovascular diseases and MI so far (summarized in Supplementary Table 3)^44-46^. However, in the APOLLO trial^44^ the mean LVEF was 52% at baseline (which is considered an incorrect target population^8^); in the PRECISE trial^45^ the LVEF was not investigated with CMR imaging; and the ATHENA trial was put on hold because of delivery related cerebrovascular events^46^. Some authors^17^ erroneously reviewed the MyStromalCell trial^47^ as a clinical trial on ADRCs for chronic MI. However, in the MyStromalCell trial ASCs stimulated with VEGF-A_165_ were used^47^. Collectively, these data clearly demonstrate that at least for the delivery of ADRCs in MI there is currently no scientific basis for dampening enthusiasm and expectations in connection with potential therapeutic success (obviously considering potential issues with certain delivery routes which may not be optimal).

Some authors^6^ proposed to focus further research into cell-based therapies for MI on the combination of the optimal cell type, dose, delivery route and timing. However, such approaches can only consider those types of cells and delivery routes that are known at a certain time – and may immediately be outdated by the description of a novel type of cell (or, in case of ADRCs, a novel method for isolating the cells) and/or another delivery route that perform better than the existing ones. This may be illustrated by the results shown in Fig. 12. In this regard it should be mentioned that at present it remains unknown to which extent the favorable outcome of the present study was at least in part due to the novel cell delivery route. Similar delivery routes on porcine animal models of MI were described in the literature^48,49^. However, the results of these studies cannot be compared to the results presented here because in both studies^48,49^, the LAD was occluded for only 45 min, cells were delivered at six days after experimental induction of MI, only short-term effects of cell retention at one or 24 h after cell delivery were investigated, and therapeutic outcome was not assessed with CMR imaging. In any case, from a clinical point of view the optimal delivery of less efficient but safer cells (such as UA-ADRCs compared to other types of cells) may be preferable compared with suboptimal delivery of more efficient, but less safe cells (such as genetically modified ASCs).

With respect to cell-free therapies for MI a recent study should be mentioned in which human ASCs of passage 3 or conditioned medium (CM) collected from the same amount of ASCs (i.e., a cell-free therapy) were intramyocardially delivered into the heart of mice immediately after ligation of the LAD artery^50^ (note that CM can in principle not be collected from ADRCs). The therapeutic efficacy of ASCs was moderately superior to CM, both ASCs and CM significantly reduced cardiomyocyte apoptosis in the infarct border zone, and ASCs enhanced angiogenesis in the infarct border zone to a stronger degree than CM^50^. These results are in line with the finding of another study that in vitro, the direct contact between cardiomyocytes and ASCs (in addition to soluble signaling molecules) is obligatory in relaying the external cues of the microenvironment controlling the differentiation of ADSCs^51^. Besides this, these results support the findings of the present study, although the experimental design of the present study did not allow to determine to what extent the increased angiogenesis (represented by a statistically higher mean microvessel density in the infarct border zone after delivery of UA-ADRCs compared to the delivery of saline) was caused by transdifferentiation of ADRCs into cells of vessel walls or by paracrine effects mediated by the delivered ADRCs. Future research should directly compare the outcome of treatment of experimentally induced MI with ADRCs, ASCs and CM in relevant animal models (i.e., using the same instrumentation and standard of care as in humans, and using CMR imaging for assessing the therapeutic outcome).

The present study has a number of limitations. First, only a single combination of cell type, dose, delivery route and timing was tested on a pre-known location of MI with uniform infarct size. However, this has been a common feature of all feasibility studies on cell-based therapies for MI, and the therapeutic outcome reported here compared with results of other studies on porcine animal models of cell-based therapies for MI in which functional outcome was assessed with CMR imaging justifies the chosen procedure. Second, the follow-up period after delivery of UA-ADRC was only six weeks. However, this was longer than both the median (4 weeks) and the mean (5.75 weeks) follow-up period (range, 2-12) of the 16 pre-clinical studies on cell-based therapy for MI using porcine and sheep animal models summarized in Supplementary Table 1. Third, the exact molecular and cellular effects that were the basis of the therapeutic outcome could not be identified (mostly because UA-ADRCs can in principle not be labeled, and no conditioned medium can be collected from UA-ADRCs). This will require direct comparison of the outcome of treatment of experimentally induced MI with ADRCs, ASCs and CM in relevant animal models, and that such studies quantify microvessels and total numbers of cardiomyocytes using state-of-the-art quantitative histologic methods^52-54^. In any case, findings from other studies in which different types of cells were delivered and that reported inhibited cardiac fibroblast growth and reduced collagen expression, beneficial effects on the ratio between matrix metalloproteinases and tissue inhibitors of metalloproteinases, as well as limited local inflammation^55-58^ may also apply after delivery of UA-ADRCs in MI. Furthermore, based on the results obtained with labeled ASCs in the present study (shown in Figs. 8-10) it is reasonable to hypothesize that at least a portion of the delivered ADRCs proliferated and integrated into the wall of vascular structures in the left ventricular border zone of the MI. Fourth, we are not aware of any comparable study in a large animal model of cell-based therapy for experimentally induced chronic MI in which application of cells was performed using the same instrumentation and standard of care as in humans and the functional outcome was assessed with CMR imaging. Nevertheless, in the present study hemodynamic endpoints were investigated that only can support but not replace relevant clinical endpoints including quality of life assessment, number of hospitalizations, 6 min walk tests, and death over several years^8^. In conclusion, the present study suggests that the delivery of UA-ADRCs at four weeks after MI is safe and effective, leading to a significant increase in the LVEF and a significant reduction in the relative amount of scar volume of the left ventricular wall, without adverse effects. Accordingly, the results of the present study justify the evaluation of a new combination of ADRCs (including the isolation procedure used), dose, delivery route and timing presented here in future clinical trials according to the strict criteria established by Madonna et al.^8^, which includes the use of CMR imaging and clinically relevant endpoints.

## METHODS

### Animals

The present study was performed on n=32 pigs. All experiments were performed in accordance with the guidelines published in NIH publication No. 86-23 titled Guide for the Care and Use of Laboratory Animals (revised 1985) and under a protocol reviewed and approved by the Institutional Animal Care and Use Committee at Houston Methodist Hospital (Houston, TX, USA) (AUP-0910-0019).

For the main experiment, twenty-five animals were randomly assigned to respectively treatment with unmodified, autologous adipose-derived regenerative cells (UA-ADRCs) (n=13; Group 1) or sham-treatment with saline (n=12; Group 2). Eight of these animals could not be included in the final analysis, for several reasons: four animals died during MI induction at time point T0; one animal had to be euthanized due to a musculoskeletal injury before the end of the study; one animal died during anesthesia at time point T0; in one animal the injection of UA-ADRCs deviated from the protocol; and one animal had a pre-existing cardiac abnormality. As a result, the final analysis of the main experiment was performed on data from nine animals in Group 1 and eight animals in Group 2.

An additional experiment was performed on five animals that were randomly assigned to respectively treatment with autologous, adipose-derived stem cells that were labeled with enhanced green fluorescent protein (eGFP^+^ A-ARCs) (n=3; Group 3) or sham-treatment with saline (n=2; Group 4). One animal in Group 3 had to be euthanized one week before the planned cell delivery.

For a third experiment, one animal received an injection of A-ASCs labeled with iron oxide particles into the left and right renal artery, and another animal an injection of A-ASCs labeled with iron oxide particles into the left anterior descending (LAD) vein (Group 5). The corresponding methods are described in Supplementary Figs 3 and 4.

### Induction of myocardial infarction (MI)

Myocardial infarction was induced in all animals in Groups 1-4 at T0 (c.f. Table 1). As previously described^38^, anti-platelet therapy was administered orally, consisting of 325 mg Acetylsalicylic Acid (Aspirin; Bayer, Leverkusen, Germany) two days before T0 (i.e, on day T0-2) as well as on T0-1, and Clopidogrel (Plavix; Sanofi-Aventis Pharma, Paris, France) with a loading dose of 300 mg on day T0-2 and 75 mg on days T0-1 and T0). In addition, the animals received the anti-arrhythmic, Bisoprolol (Concor; Merck, Darmstadt, Germany) from T0-2 to T0+5 (1.25 mg per day orally).

After establishing vessel access, anti-coagulant therapy was administered intravenously as follows: Acetylsalicylic Acid 500 mg (Aspisol, Bayer, Leverkusen, Germany), Enoxaparin 1 mg/kg bolus, then 0.5 mg/kg every 4h (Lovenox; Sanofi-Aventis Pharma), and Eptifibatide prior to balloon occlusion in two 180 µg/kg boluses, 10 min apart, followed by a 2 µg/kg/min infusion during balloon occlusion (Integrilin, SP Europe, Bruxelles, Belgium).

A baseline coronary angiography (CA) (Fig. 1a) and a ventriculography in right and left anterior oblique views (RAO/LAO VG) was performed using a fluoroscopy C-arm (Axiom ArtisTM; Siemens, Erlangen, Germany). Then, a coronary angioplasty balloon (length 8-12 mm, diameter 3.0-3.5 mm; Maverick OTW; Boston Scientific, Marlborough, USA) was directed over a 0.014” guide wire (Choice Floppy; Boston Scientific) into the mid left anterior descending (LAD) artery and inflated for three hours at the minimal pressure (typically 2 atm) required for a complete occlusion (Fig 1b). After three hours, the balloon was deflated and a post-MI CA was performed to insure vessel patency (Fig 1c). Catheters and sheaths were removed and the animal was taken cared for as previously described^38^. Enoxaparin 1 mg/kg (Lovenox) was subcutaneously administered at the end of the MI induction procedure.

### Isolation of adipose-derived regenerative cells (ADRCs)

Cells were isolated either during CA and RAO/LAO VG at the time of MI induction (i.e., at T0) (Groups 3 and 4) or during CA and LAO/RAO VG four weeks after T0 (i.e., at time point T1) (Groups 1 and 2), respectively (c.f. Table 1). Following incision with a scalpel, 12-25 g of subcutaneous adipose tissue was harvested from the nuchal region of each pig. The tissue was divided into aliquots of about 6 to 10 g each. Then, each aliquot was processed using the Transpose RT system (InGeneron) for isolating ADRCs from adipose tissue. To this end, each aliquot was minced and incubated together with Matrase enzyme blend^59,60^ (InGeneron) for 1 h under agitation in the processing unit at 39° C according to the manufacturer’s instructions for use. For the preparations of UA-ADRCs of the animals in Groups 1 and 2, cell counting, colony forming unit assay and flow cytometry was performed.

### Counting and evaluation of cell viability

Cells were stained with fluorescent nucleic acid stain (SYTO13; Life Technologies, Grand Island, NY, USA) following manufacturer’s instructions, and then counted using a hemocytometer under an Eclipse Ti-E inverted fluorescence microscope (Nikon Corporation, Tokyo, Japan) using a PlanFluor 10× objective (numerical aperture [NA] = 0.3) (Nikon).

Viability of UA-ADRCs was determined by preparing a 3:1 dilution of the cell suspension in 0.4% Trypan Blue solution. Nonviable cells were counted using a hemocytometer under the same microscope operated in fluorescence mode, and were correlated to the number of viable nucleated cells.

### Colony-forming unit (CFU) assay

For each preparation of UA-ADRCs (Groups 1 and 2), freshly isolated nucleated cells were plated at a density of 100,000 cells per 35-mm-diameter well in triplicate, and incubated for 14 days in standard growth media as previously described^61^. Media were changed twice weekly. Afterwards the cells were fixed with 4% formalin, stained with hematoxylin for 10 minutes and washed with phosphate-buffered saline (PBS). Photomicrographs were taken from five randomly chosen fields-of-view per well with a DS-Fi1 CCD color camera (2560 x 1920 pixels; Nikon) attached to an Eclipse Ti-E inverted microscope (Nikon) and NIS-Elements AR software (version 4.13; Nikon), using a Plan EPI 2.5× objective (NA = 0.075) (Nikon). A CFU was defined as a cluster containing at least 10 fibroblast-like fusiform cells^60^. Two different experienced investigators individually counted all visible colonies.

### Flow cytometric analysis

Fresh ADRCs were cultured for 24 h, incubated with antibodies for 30 min, washed, re-suspended in 1 ml PBS with 10% fetal bovine serum (FBS) and 1% sodium azide, and directly analyzed by flow cytometry using a BD FACSAria Fusion device (BD Bioscience, San Jose, CA, USA). Antibodies against porcine CD29 (antibodies-online, Aachen, Germany), CD44 (Abcam, Cambridge, MA, USA), NG2 (Abcam), Oct4 (Novus Biologicals, Littleton, CO, USA), CD31 (Gentex, Irvine, CA, USA), CD45 (Abcam), Nestin (Santa Cruz Biotechnology, Dallas, TX, USA), CD146 (Gentex, Irvine, CA, USA) and CD117 (eBioscience, San Diego, CA, USA) were used.

### Labeling of cells with enhanced green fluorescent protein

Cells isolated from subcutaneous adipose tissue of animals in Groups 3 and 4 were expanded in cell culture for 5-7 days. At passage 3 the cells were simultaneously transfected (using FuGENE 6 Transfection Reagent; Promega Coporation, Madison, WI, USA) with plasmids encoding eGFP fused to the nuclear localization signal H2B and other plasmids containing PiggyBac Transposase (System Bioscience, Mountain View, CA, USA) which was transiently expressed in order to integrate the eGFP cargo into the genome. After transfection, eGFP positive cells were selected for 14 days in complete growth media containing 400 ng/ml G418 (Life Technologies). Then, cells were separated using fluorescence-activated cell sorting (FACS) using the aforementioned BD FACSAria Fusion device (BD Bioscience). Sorted cells (with >95% of the cells expressing eGFP) were expanded for additional 5-7 days in cell culture. On the day of delivery, eGFP^+^ cells were trypsinized for 5 min at 37° C, washed twice with PBS, centrifuged at 600 g for 10 min, passed through a 70 µm cell strainer (Falcon, Corning, NY, USA) to avoid cell clumping, and suspended in 10 ml sterile saline solution for delivery (B. Braun Medical Inc., Bethlehem, PA, USA) (on average 10×10^6^ cells per animal). Cells were counted as described above; expression of eGFP was confirmed by fluorescence microscopy during cell counting.

### Delivery of cells or saline as control

Either UA-ADRCs (Group 1), eGFP^+^ A-ASCs (Group 3) or saline (Groups 2 and 4) were delivered during CA through a standard over-the-wire balloon catheter that was placed and inflated in the LAD vein that corresponded to the previous LAD artery occlusion site at T1 (i.e., four weeks after T0) (Fig. 1d and Table 1). To this end, a 6F Amplatz right 1 guide catheter (Mach 1; Boston Scientific) was advanced over a 0.035’’ wire (J-tip Starter; Boston Scientific) through the jugular vein into the coronary sinus (CS). Once in place in the CS, an angioplasty balloon (length 8-12 mm; diameter 3.0-3.5 mm; Maverick OTW; Boston Scientific) was positioned over a 0.014’’ guide wire (Choice Floppy; Boston Scientific) at the site of the LAD vein. The wire was removed and a suspension of UA-ADRCs (18×10^6^ cells in 10 ml saline; Group 1), eGFP^+^ A-ASCs (on average 10×10^6^ cells in 10 ml saline; Group 3) or 10 ml saline alone (Groups 2 and 4) were delivered at a rate of ∼0.25 ml/s through the catheter’s central lumen retrogradely into the LAD vein. The angioplasty balloon was kept inflated during the entire delivery procedure and the five minutes following the delivery procedure. Operators were blinded to the group assignment.

### Cardiac magnetic resonance (CMR) imaging

All animals underwent CMR imaging directly before the delivery of UA-ADRCs or saline at T1 as well as at T2. The scans were performed using a 1.5 Tesla MRI scanner (Avanto, Siemens Medical Solutions, Inc., Erlangen, Germany).

### Steady-state free precession (SSFP) CMR imaging

Cine-CMR images were acquired using an electrocardiogram-gated steady-state free precession pulse sequence in multiple short-axis and long-axis views. Short-axis views were obtained every 1 cm from the atrioventricular ring to the apex to cover the entire left ventricle (slice thickness 6 mm, inter-slice gap 4 mm, echo time (TE) 1-1.5 ms, temporal resolution (TR) 35–45 ms, flip angle 50°-90°).

### Late gadolinium enhancement (LGE) CMR imaging

Approximately ten minutes after the administration of 0.15 to 0.20 mmol/kg of gadolinium (Magnevist, Bayer Inc. Mississauga, ON, Canada), LGE-CMR imaging was performed in slice orientations identical to cine-CMR imaging using a standard inversion recovery gradient echo pulse sequence. Manual adjustment of the time from inversion (TI) was performed in all cases in order to achieve nulling of normal viable myocardium^62^. Typical imaging parameters were matrix 256×192, slice thickness 6 mm, gap 4 mm, TI 250–350 ms, TE 2.0–2.5 ms, flip angle 30°, and parallel imaging with two-fold acceleration factor.

### Analysis of CMR images

Image analysis was performed with the cvi^42^ software (Circle Cardiovascular Imaging Inc., Calgary, AB, Canada). Endocardial and epicardial borders were traced using planimetry on stacks of short-axis cine images in end diastole and end systole. Based on these data the following variables were calculated for each animal: cardiac output, stroke volume, left ventricular ejection fraction (LVEF), end-diastolic volume (EDV), end-systolic volume (ESV) and left ventricular mass. The LVEF was calculated as the difference between EDV and ESV, divided by EDV. The left ventricular mass was calculated by the difference between the left ventricular epicardial and endocardial volumes during end systole, multiplied by the myocardial density (1.05 g/cm^3^)^63^. For quantification of scar tissue, a region of normal myocardium was independently chosen on each short axis image showing the left ventricle, and hyper-enhanced regions in each slice were located. Hyper-enhancement was identified as areas of signal intensity ≥ 5 standard deviations greater than normal myocardium^35^. Two well trained MRI physicians, blinded to the group assignment, evaluated the selected regions and graded the findings in all segments on each short axis image.

### Termination

Animals were euthanized according to the Houston Methodist Research Institute Euthanasia for Large Animals Procedure (Houston, TX, USA) after performing CA, RAO/LAO VG and CMR imaging at T2. Intravenous injection of 0.25 mg/kg Pentobarbital/Phenytoin combination was performed in conjunction with isoflurane overdose. Death was verified by absence of vital signs. Hearts and organs were removed for further histologic and immunofluorescence analysis. The animal carcasses were disposed of in accordance to standard operating procedures of the Houston Methodist Research Institute (Houston, TX, USA).

### Histologic processing of heart tissue

Hearts were harvested and fixed in 5% paraformaldehyde. The left ventricle was cut into six transversal, 1 cm-thick slices from apex to base. Then, from each heart several, approximately 1×1×1 cm large tissue samples were collected, representing the left ventricular borderzone of MI, the core region of MI, and regions of viable myocardium. Specimens were paraffin-embedded and cut into 5 µm-thick tissue sections that were mounted on glass slides and stained with Masson’s Trichrome staining or processed with fluorescence immunohistochemistry.

### Evaluation of microvessel density

The microvessel density was determined on up to four representative sections from each animal in Groups 1 and 2 showing the left ventricular borderzone of MI. Only sections that showed at least 30% of both scar tissue and viable myocardium were considered in this analysis. To this end, photomicrographs covering the entire section were taken with a DS-Fi1 CCD color camera (Nikon) attached to an Eclipse Ti-E inverted microscope (Nikon) using a PlanApo 20× objective (NA = 0.75) (Nikon). Images were analyzed using ImageJ software (U. S. National Institutes of Health, Bethesda, MD, USA). Using a defined grid of ten fields per section with an area of 0.3 mm^2^ per field, two independent, blinded evaluators determined the number of microvessels per field (microvessels with a diameter between 2 and 10 µm were counted). Microvessel density was calculated based on microvessel counts on a total of 202 fields (animals in Group 1) or 247 fields (animals in Group 2), respectively.

### Immunohistochemistry

After de-paraffinizing and rehydrating, sections were washed with PBS containing 0.3% Triton X-100 (Sigma Aldrich, St. Louis, MO, USA) and blocked with 10% casein solution (Vector Laboratories, Burlingame, CA, USA) for 30 min at room temperature. Sections were incubated overnight with diluted primary antibodies and subsequently with diluted secondary antibodies for 1 h. The following antibodies were used in the present study: Goat anti-GFP, Mouse anti-cardiac troponin T, Rabbit anti-CX43, Rabbit anti-von Willebrand factor (vWF) (all from Abcam); Mouse anti-Ki67, Alexa Fluor 647 conjugated (BD Bioscience); Rabbit anti-adiponectin, Cy5 conjugated (Biorbyt, San Francisco, CA, USA), Rabbit anti-goat-IgG secondary antibody, FITC conjugated, Goat anti rabbit-IgG secondary antibody, Cy5 conjugated, Donkey anti-mouse-IgG secondary antibody, TRITC conjugated (all from Life Technologies, Carlsbad, CA, USA), and Goat anti-chicken-IgG secondary antibody, Texas red conjugated (Thermo Scientific, Waltham, MA, USA). Counterstaining of nuclei and mounting were performed with Vectashield Antifade Mounting Medium with DAPI (Vector Laboratories).

### Photography

The photographs shown in Fig. 5 were taken with a PowerShot SD 1000 digital camera (Canon, Tokyo, Japan). The photomicrographs shown in Figs 6-11 were produced by digital photography using a DS-Fi1 CCD color camera (Nikon) (Fig. 6) or a CoolSNAP HQ2 CCD monochrome camera (1392 x 1040 pixels; Photometrics, Tucson, AZ, USA) (Figs 7-11) attached to an Eclipse Ti-E inverted microscope (Nikon) and NIS-Elements AR software (Nikon), using the following objectives (all from Nikon): PlanFluor 4× (NA = 0.13), 10× (NA = 0.3), 20× (NA = 0.45) and PlanApo 20× (NA = 0.75), 40× (NA = 0.95) and 60× (oil; NA = 1.4). Merged figures were constructed using ImageJ software (version 1.51j8; U.S. National Institutes of Health). The final figures were constructed using Corel Photo-Paint X7 and Corel Draw X7 (both versions 17.5.0.907; Corel, Ottawa, Canada). Only minor adjustments of contrast and brightness were made using Corel Photo-Paint, without altering the appearance of the original materials.

### Statistical analysis

For all investigated variables (cardiac output, stroke volume, left ventricular ejection fraction, end-diastolic volume, end-systolic volume, left ventricular mass, relative amount of scar tissue and microvessel density) mean and standard error of the mean were calculated at both investigated time points T1 and T2 (microvessel density was investigated only at T2) separately for the animals in Group 1 and the animals in Group 2. This was done after having tested with with the Shapiro-Wilk normality test whether the values came from a Gaussian distribution. Except of the microvessel density, comparisons between groups were performed with repeated measures two-way analysis of variance followed by post hoc Bonferroni tests for pairwise comparisons. For microvessel density, comparisons between groups were performed with the unpaired two-tailed Student’s t-test. In all analyses an effect was considered statistically significant if its associated p value was smaller than 0.05. Calculations were performed using GraphPad Prism (version 7.0 for Windows, GraphPad software, San Diego, CA, USA).

### Data availability

All data supporting the results of these studies are available within the paper, the associated Supplementary Materials, or from the authors upon reasonable request.

## Supporting Information

## SUPPLEMENTARY TABLES

**Supplementary Table 1.**
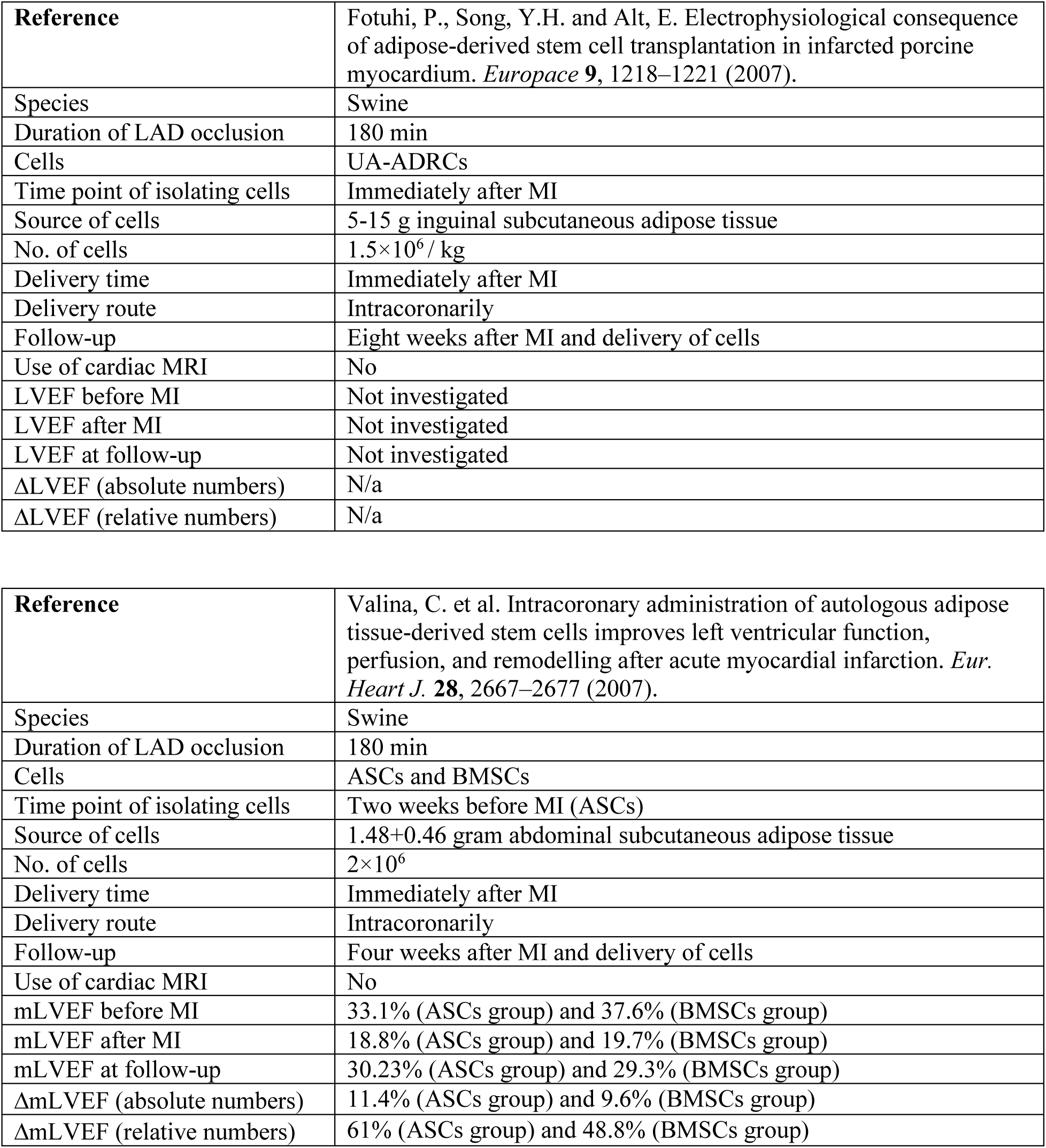
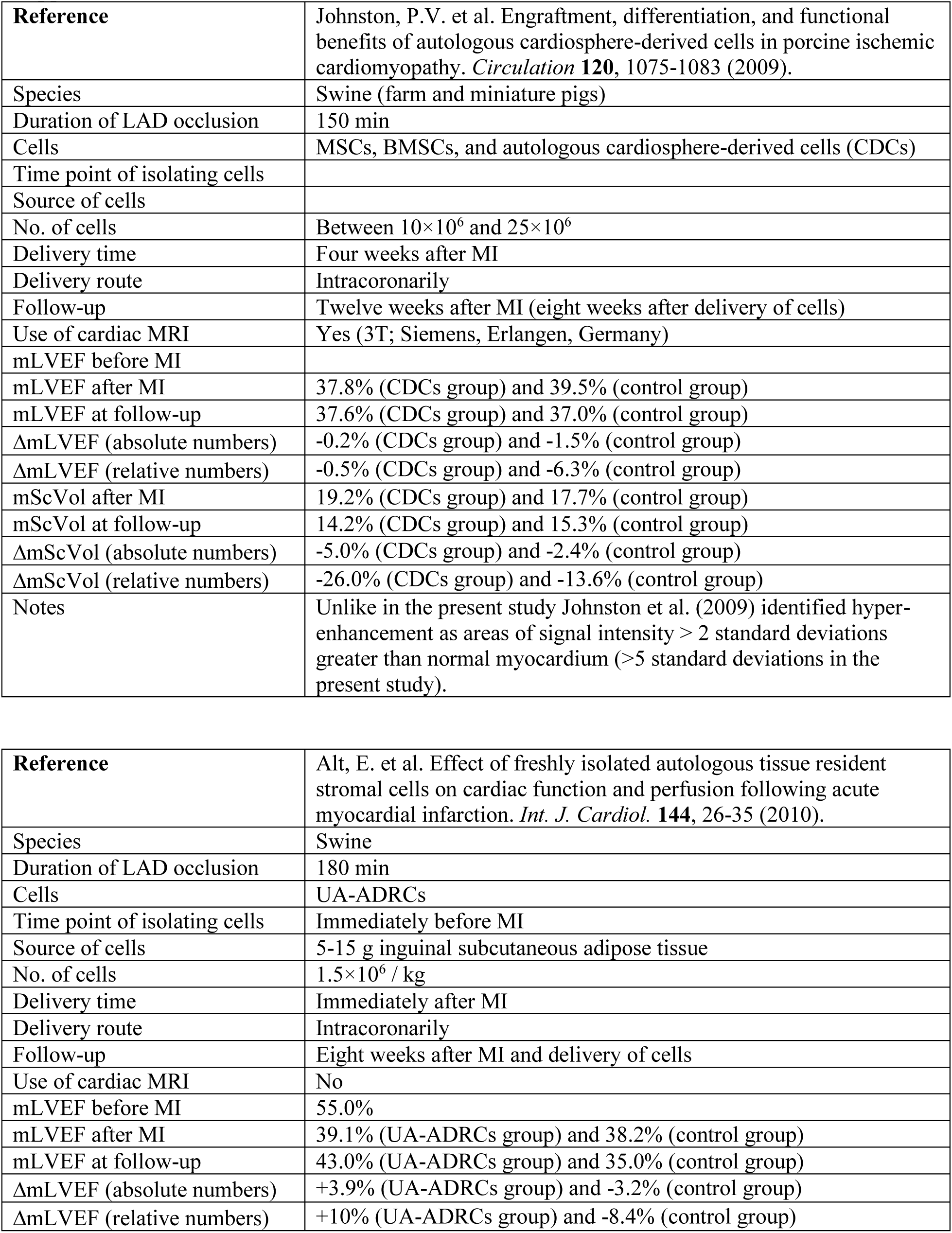
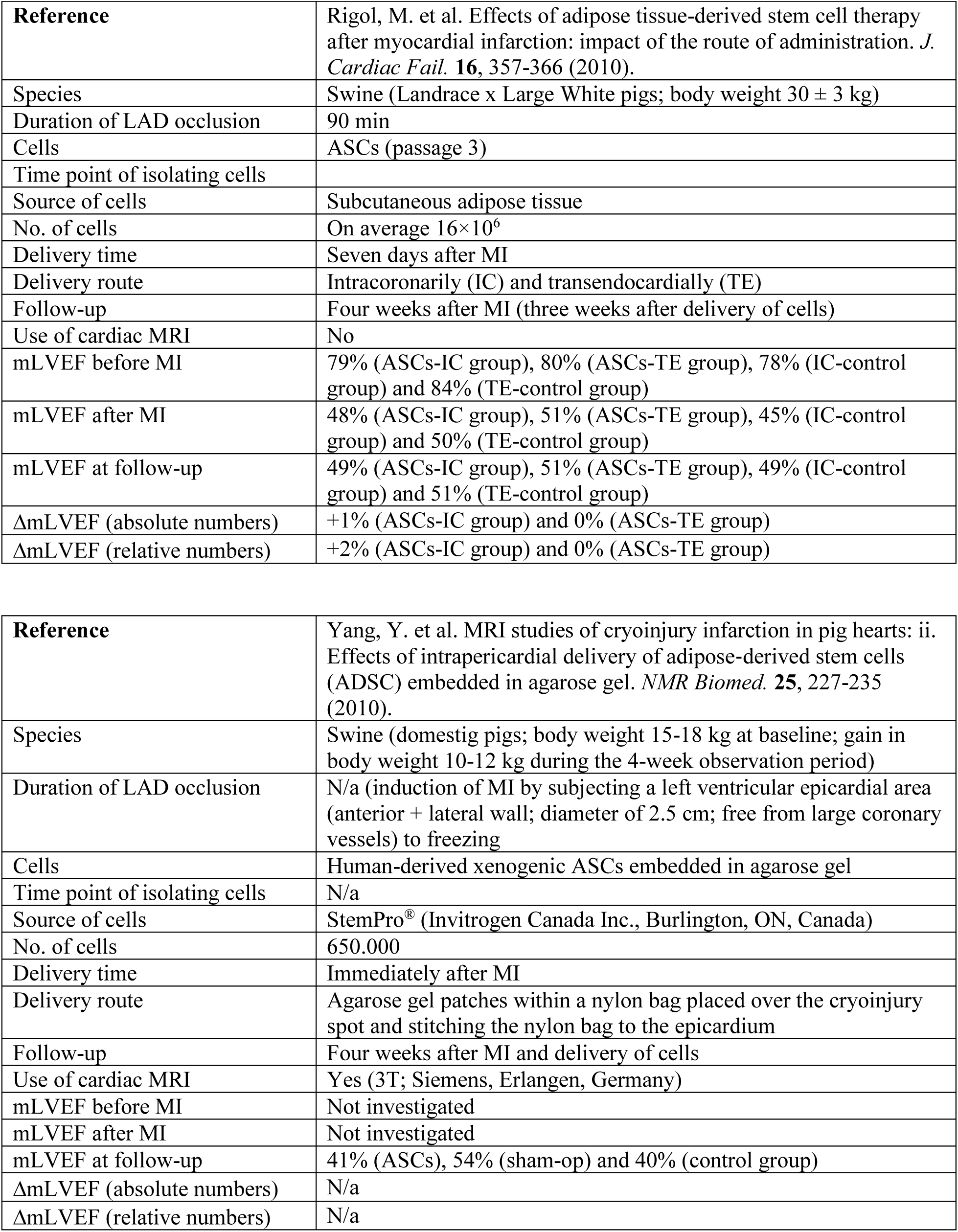
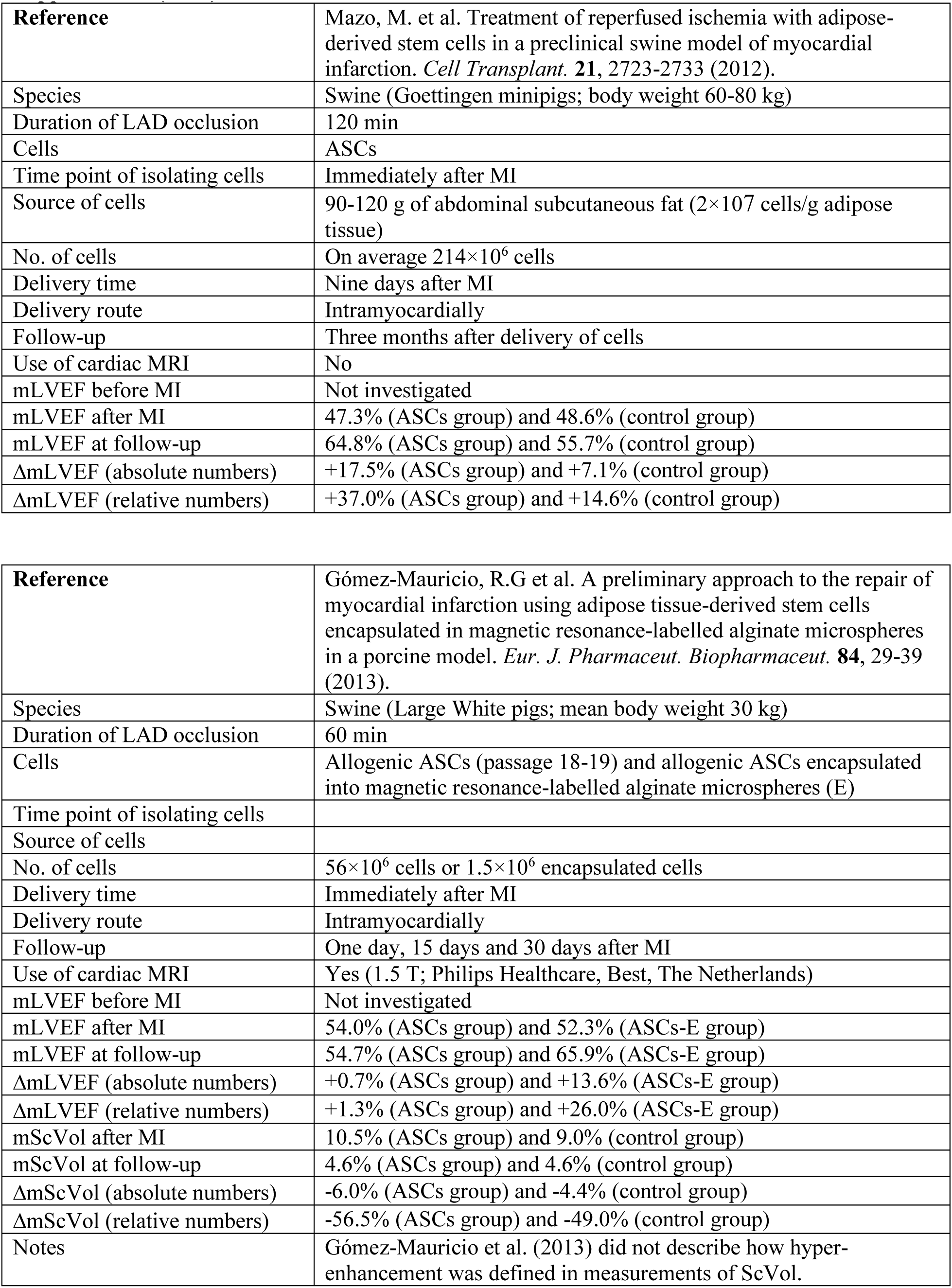
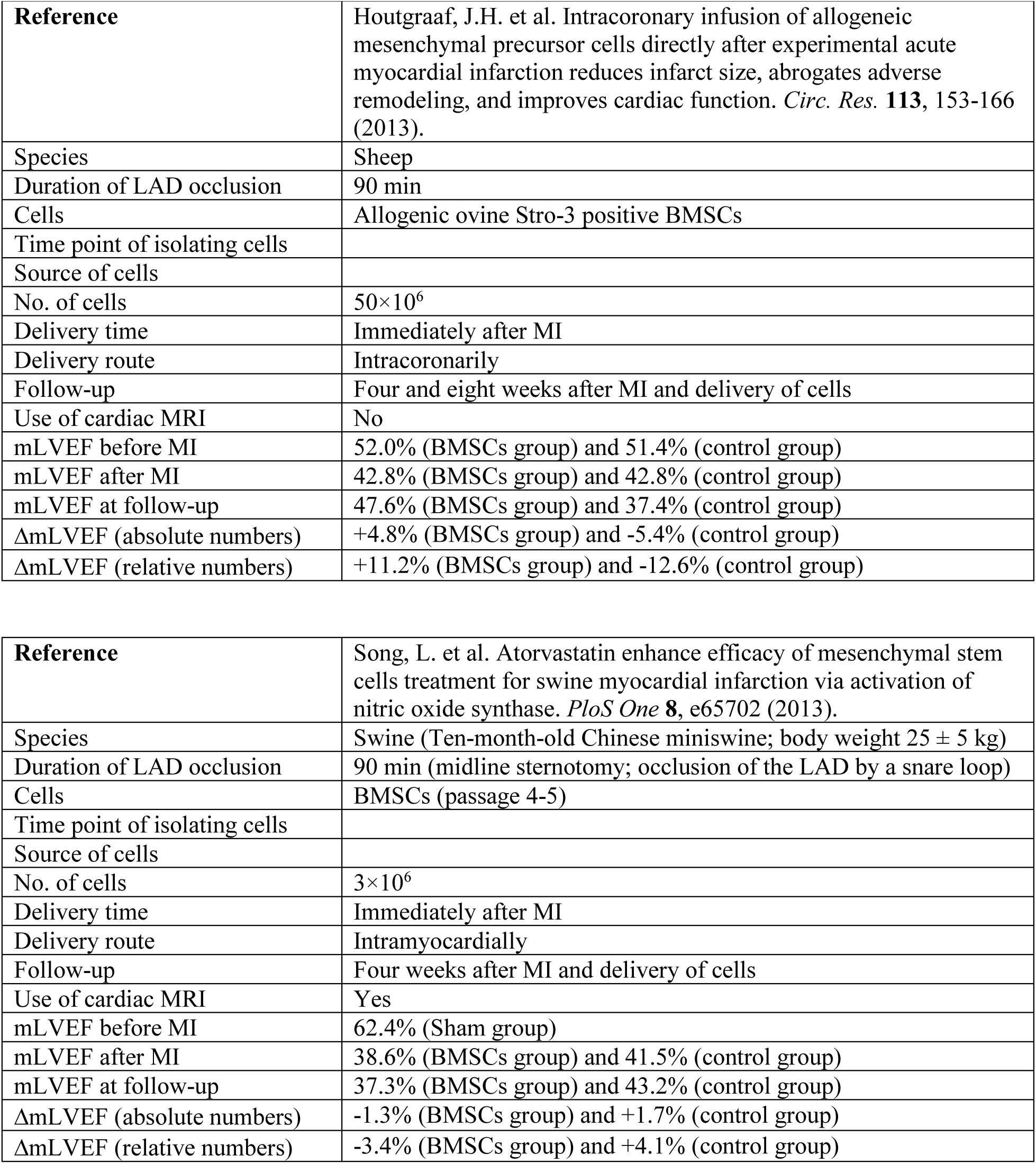
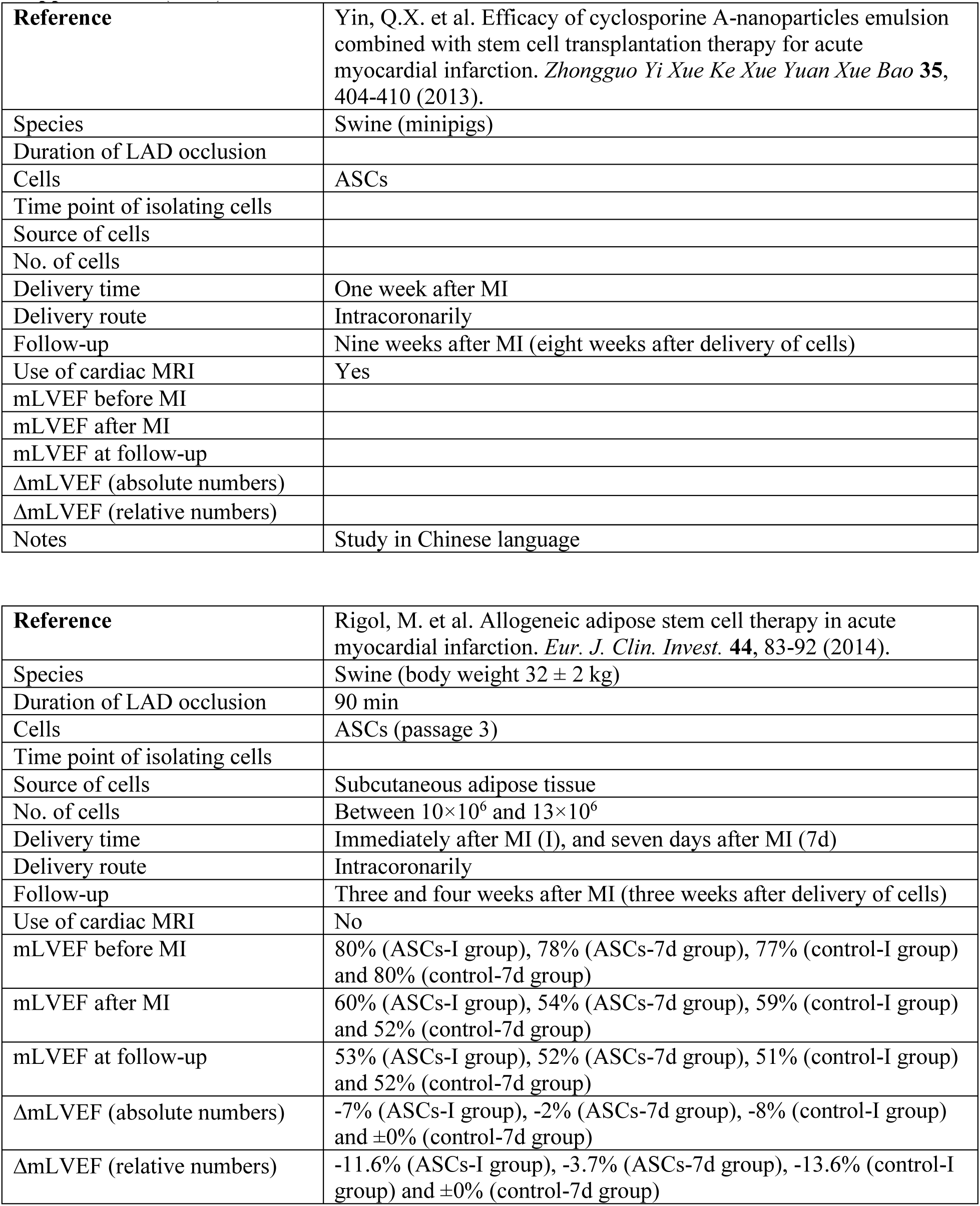
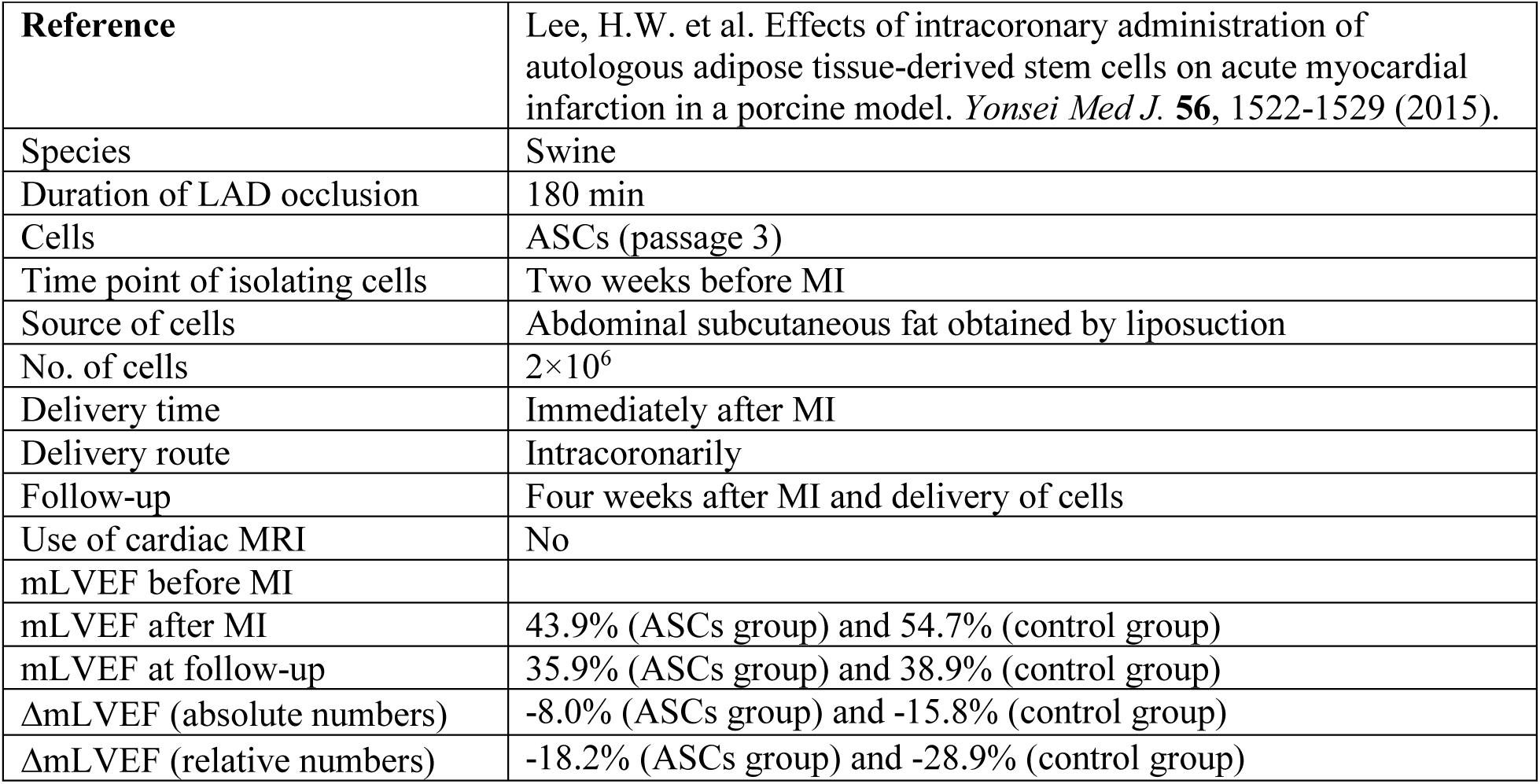
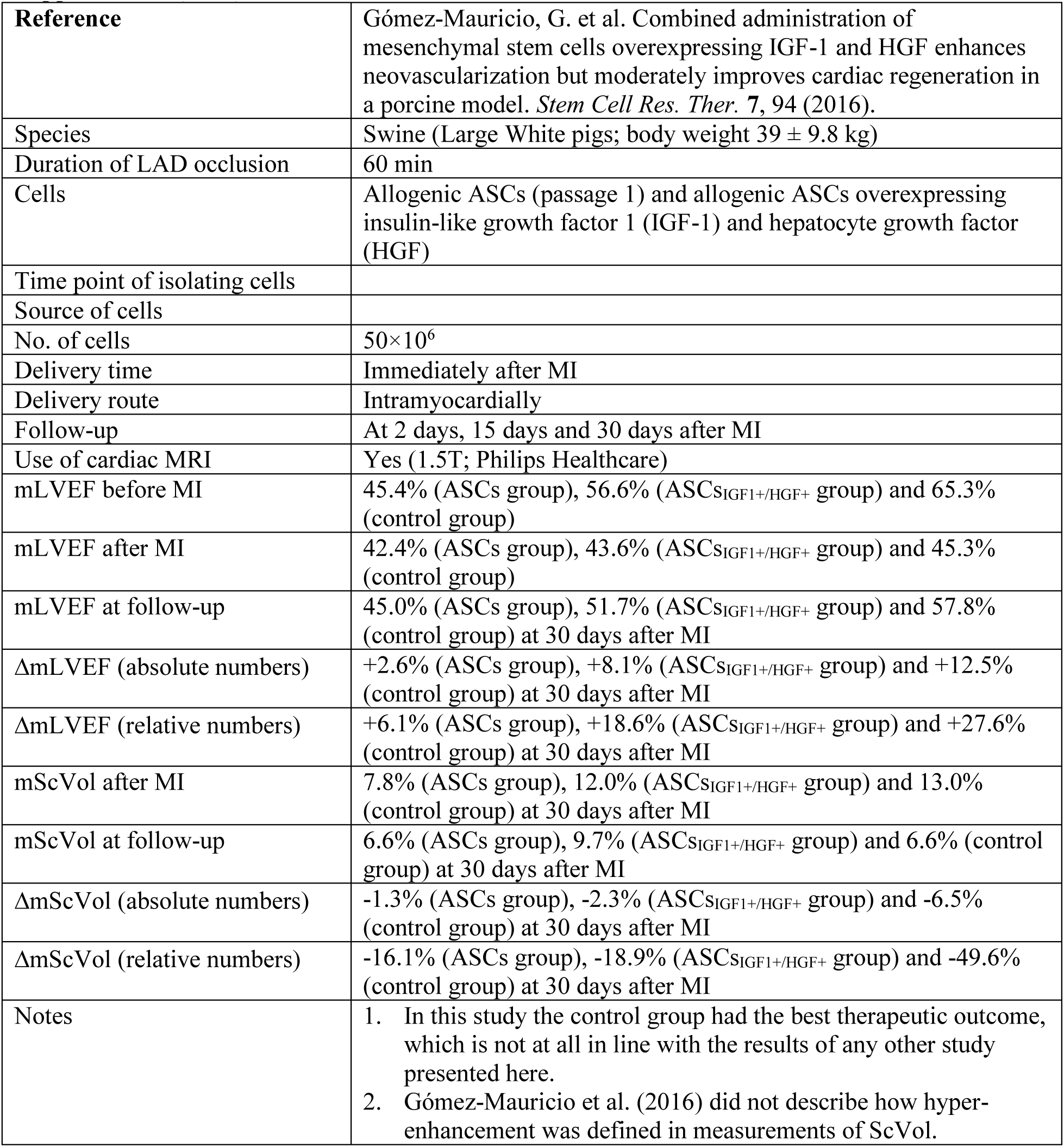
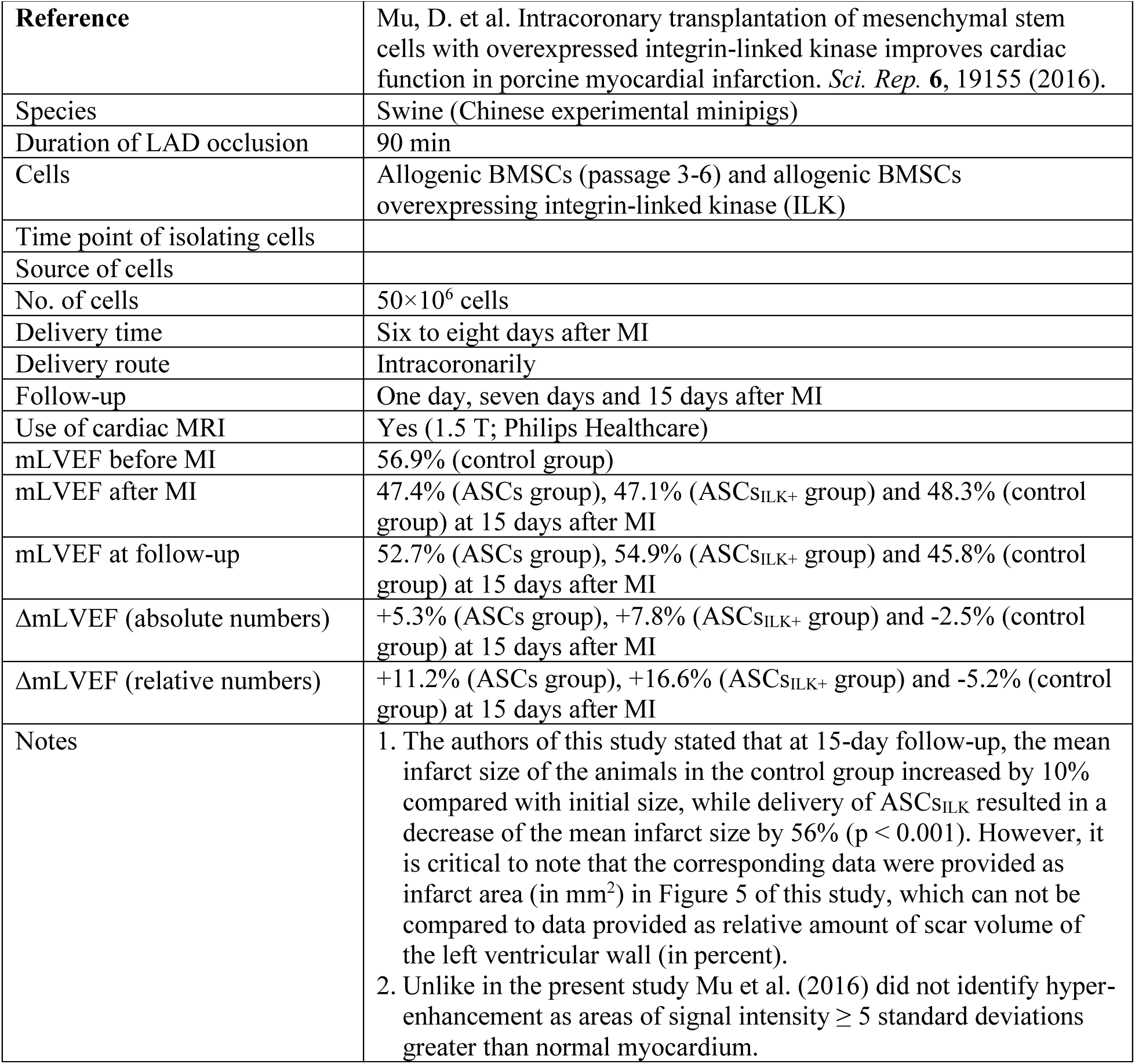
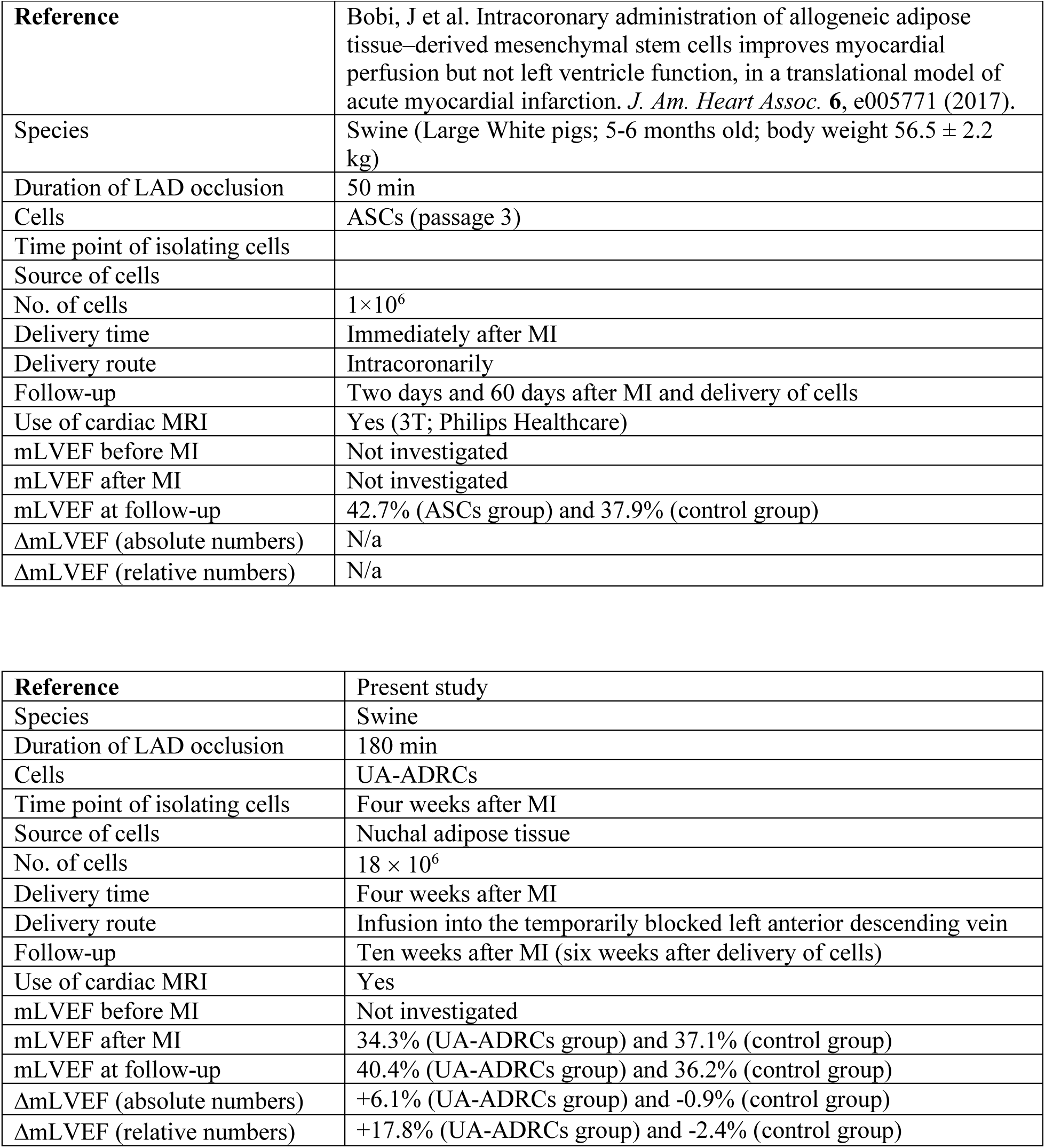
Details of studies that addressed cell therapy for experimentally induced myocardial infarction using large animal models (pigs, sheep). Abbreviations: mLVEF, mean left ventricular ejection fraction; MI, myocardial infarction; mScVol, mean relative amount of scar volume of the left ventricular wall.

**Supplementary Table 2.** Mean and standard error of the mean of functional (CO, SV, LVEF, EDV, ESV and HR) and structural alterations (M_LV_ and ScVol) of the porcine heart at ten weeks (i.e., at time point T2) after experimental occlusion of the left anterior descending (LAD) artery for three hours at time point T0, followed by the delivery of respectively 18 × 10^6^ unmodified, autologous adipose-derived regenerative cells (UA-ADRCs) (n=9) or saline (Control) (n=8) into the balloon-blocked LAD vein (matching the initial LAD artery occlusion site) at four weeks after T0 (i.e., at time point T1). Abbreviations: P, parameter; CO, cardiac output (l/min); SV, stroke volume (ml); LVEF, left ventricular ejection fraction (%); EDV, end diastolic volume (ml); ESV, end systolic volume (ml); HR, heart rate (min^-1^); M_LV_, left ventricular mass (g); ScVol, relative amount of scar volume of the left ventricular wall (%); D, difference; B, results of group-specific Bonferroni’s multiple comparison tests of the difference in the group means between T1 and T2; *, p<0.05; **, p<0.01; ***, p<0.001.

| P | Group 1 (delivery of UA-ADRCs) |  |  |  |  |  | Group 2 (delivery of saline) |  |  |  |  |  |
| --- | --- | --- | --- | --- | --- | --- | --- | --- | --- | --- | --- | --- |
|  | T1 |  | T2 |  | D | S | T1 |  | T2 |  | D | S |
|  | Mean | SEM | Mean | SEM |  |  | Mean | SEM | Mean | SEM |  |  |
| CO | 2,73 | 0,22 | 3,78 | 0,23 | +38% | ** | 3,39 | 0,30 | 3,63 | 0,32 | +7% | - |
| SV | 31,4 | 2,02 | 44,3 | 2,39 | +41% | *** | 34,6 | 2,99 | 38,0 | 3,43 | +10% | - |
| LVEF | 34,3 | 2,89 | 40,4 | 2,64 | +18% | * | 37,8 | 2,57 | 36,2 | 2,45 | -4% | - |
| EDV | 93,6 | 5,07 | 111,9 | 6,24 | +20% | *** | 92,6 | 6,45 | 106,1 | 7,17 | +15% | ** |
| ESV | 62,1 | 5,28 | 67,6 | 6,49 | +9% | - | 58,0 | 5,24 | 68,1 | 6,03 | +17% | * |
| HR | 87,0 | 2,93 | 85,4 | 4,19 | -2% | - | 97,4 | 3,96 | 95,5 | 3,61 | -2% | - |
| $M_{LV}$ | 55,3 | 4,99 | 71,3 | 4,49 | +29% | *** | 63,2 | 3,35 | 68,4 | 3,97 | +8% | - |
| ScVol | 20,9 | 2,29 | 16,6 | 1,19 | -21% | * | 17,6 | 1,38 | 22,7 | 1,76 | +29% | * |

**Supplementary Table 3.**
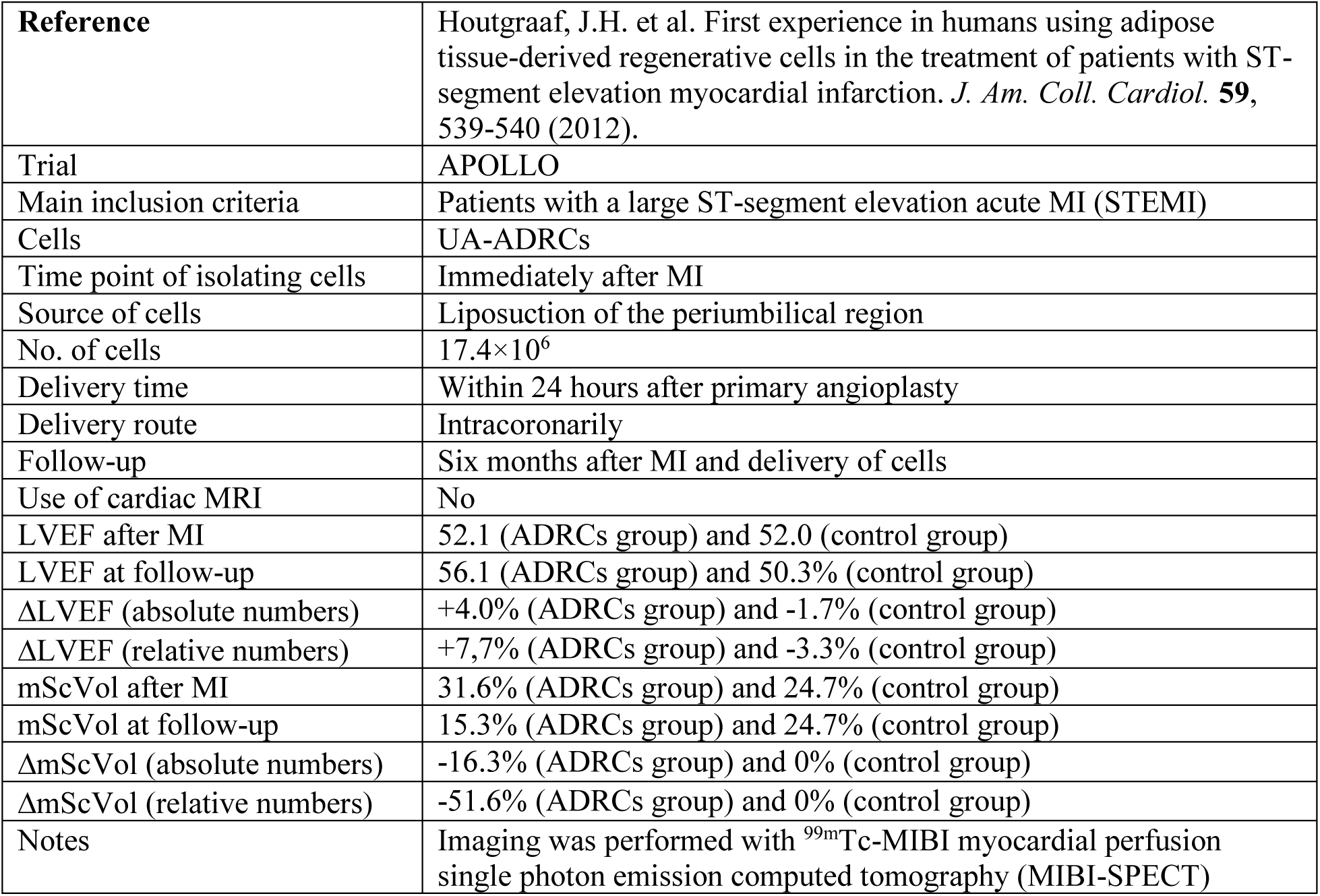
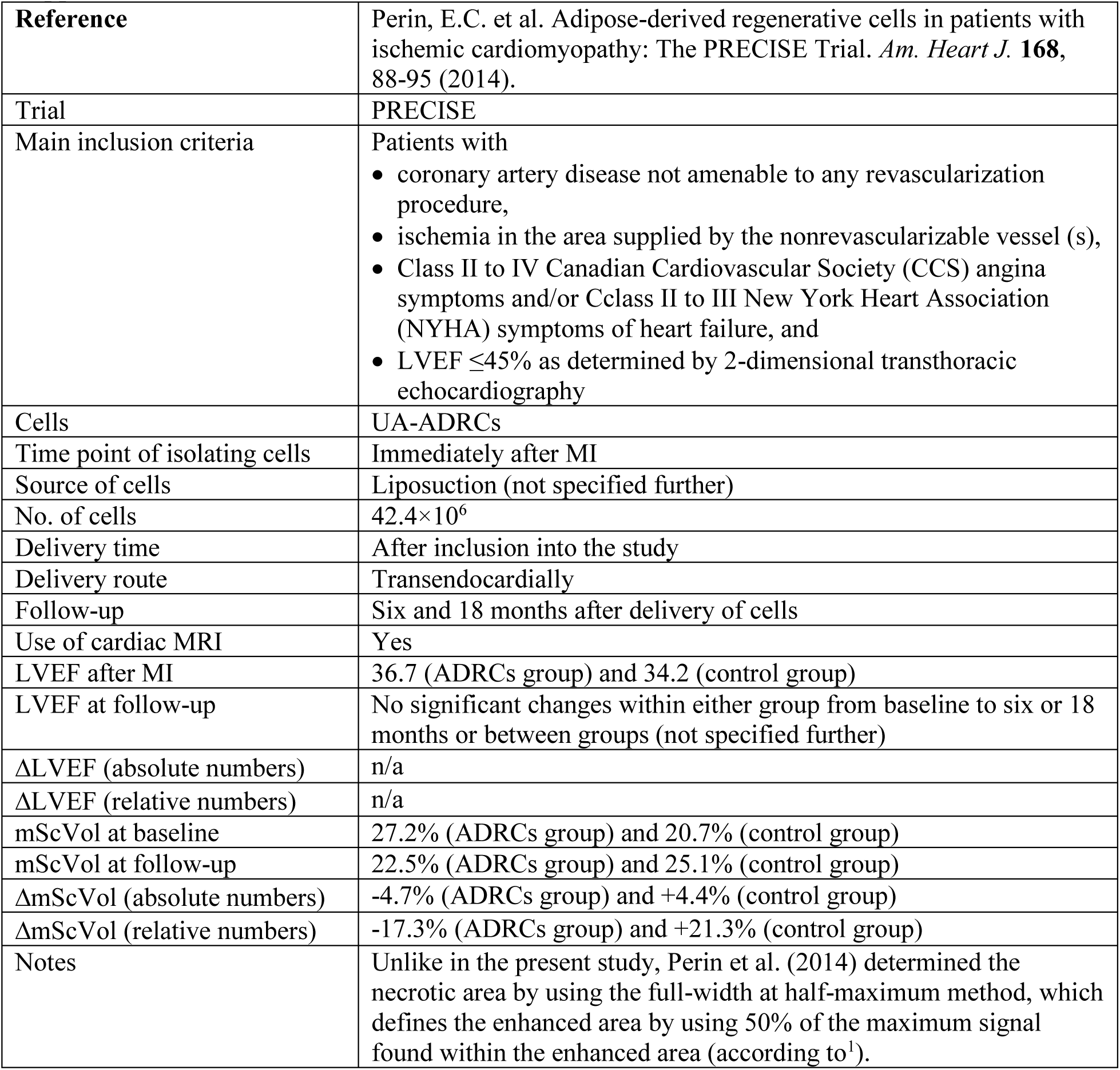
Details of studies that used adipose-derived regenerative cells for treating patients with MI. Abbreviations: mLVEF, mean left ventricular ejection fraction; MI, myocardial infarction; mScVol, mean relative amount of scar volume of the left ventricular wall.

## SUPPLEMENTARY FIGURES

**Supplementary Fig. 1.**
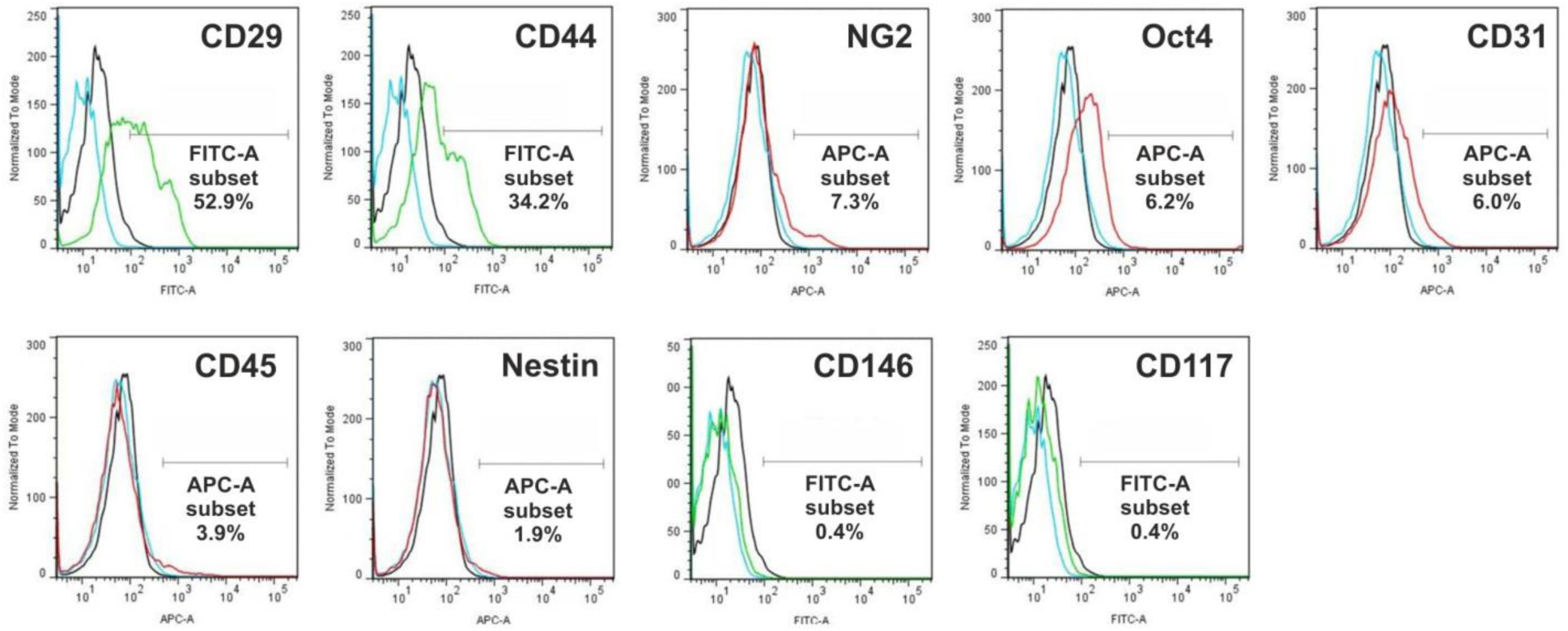
Immunophenotypic analysis of cell surface markers of freshly isolated, unmodified, autologous adipose-derived regenerative cells from an animal in Group 1 (c.f. Table 1 in the main text) using flow cytometry. The cells were stained with monoclonal antibodies for CD29, CD44, NG2, Oct4, CD31, CD45, Nestin, CD146 and CD117 at passage 0. Flow cytometric histographs are representative of triplicate experiments.

**Supplementary Fig. 2.**
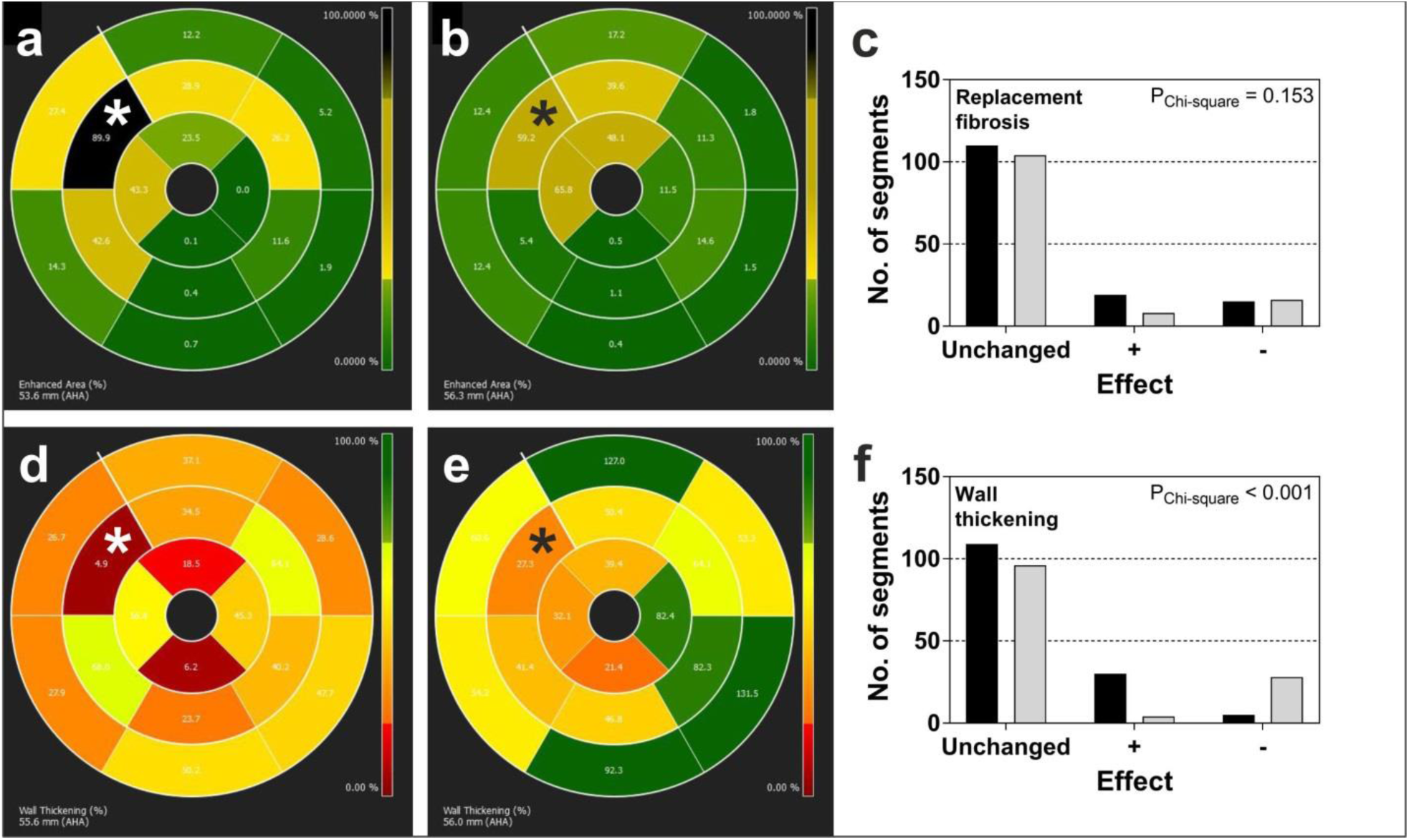
Analysis of regional replacement fibrosis (investigated with late gadolinium enhancement cardiac magnetic resonance [CMR] imaging) (**a**-**c**) and regional wall thickening (investigated with steady-state free precession CMR imaging) (**d**-**f**) of the porcine heart at ten weeks (i.e., at time point T2) after experimental occlusion of the left anterior descending (LAD) artery for three hours at time point T0, followed by the delivery of respectively unmodified, autologous adipose-derived regenerative cells (n=9) (closed bars in **c**,**f**) or saline as control (n=8) (gray bars in **c**,**f**) into the balloon-blocked LAD vein (matching the initial LAD occlusion site) at four weeks after T0 (i.e., at time point T1). Details of CMR imaging are provided in the main text. Image analysis was performed with the cvi^42®^ software (Circle Cardiovascular Imaging Inc.). For the analyses shown here, the left ventricle was divided into 17 segments as recommended by the American Heart Association (AHA)^2^. Except of the apical segment (no. 17) that cannot be investigated on short axis, transversal images through the mid left ventricle of a porcine heart as applied in this study (c.f. Fig. 2 in the main text) all segments were analyzed, yielding a total of 9×16 (Group 1) + 8×16 (Group 2) = 272 analyzed segments. The analysis of regional replacement fibrosis was performed as follows: (i) The signal intensity (SI) of a given pixel was considered hyper-enhanced when its SI was ≥ 5 standard deviations greater than the SI of normal myocardium (SI > 5 SD); otherwise its SI was considered normal^3^. (ii) For each segment the relative number of pixels with hyper-enhanced SI was calculated (NPrel_SI>5SD_). These calculations were seperately performed for T1 and T2, resulting in segment-specific data NPrel_SI>5SD_-T1 and NPrel_SI>5SD_-T2. (iii) These data were visualized using color coded, AHA 17-segment bullseye plots as shown in **a**,**b** (green: segments with viable myocardium; black: segments that showed complete fibrosis, i.e., NPrel_SI>5SD_ > 90%). (iv) For all segments of a given animal, the difference between NPrel_SI>5SD_-T1 and NPrel_SI>5SD_-T2 was calculated as ΔNPrel_SI>5SD_ = NPrel_SI>5SD_-T1 -NPrel_SI>5SD_-T2. (v) Mean and standard deviation of all 272 ΔNPrel_SI>5SD_ data were calculated. (vi) All segments with ΔNPrel_SI>5SD_ > (mean + 1 SD) were classified as improved, segments with ΔNPrel_SI>5SD_ < (mean - 1 SD) as deteriorated, and all other segments as unchanged. The analysis of regional wall thickening was performed as follows: (i) For each segment, the relative wall thickness was calculated as the difference between the end-systolic and end-diastolic wall thickness, divided by the end-systolic wall thickness (WT_rel_). These calculations were seperately performed for T1 and T2, resulting in segment-specific data WT_rel-T1_ and WT_rel-T2_. (ii) These data were also visualized using color coded, AHA 17-segment bullseye plots as shown in **c**,**d** (green: segments with viable myocardium and regular wall motion between systole and diastole; dark red: segments that showed completely akinetic myocardium). (iii) For all segments of a given animal, the difference between WT_rel-T1_ and WT_rel-T2_ was calculated as ΔWT_rel_ = WT_rel-T2_ - WT_rel-T1_. (iv) Mean and standard deviation of all 272 ΔWT_rel_ data were calculated. (iv) All segments with ΔWT > (mean + 1 SD) were classified as improved, segments with ΔWT < (mean - 1 SD) as deteriorated, and all other segments as unchanged. Statistical analysis using the Chi-square test was performed with GraphPad Prism (version 5; GraphPad, San Diego, USA). **a**,**b** Representative examples of color coded, AHA 17-segment bullseye plots, showing the results of the analysis of regional replacement fibrosis of an animal in Group 1 (internal no. 12056) at T1 (**a**) and T2 (**b**). The asterisks indicate a segment that improved between T1 and T2. **c** Number of segments with unchanged, improved (+) and deteriorated (-) signal intensity in the analysis of regional replacement fibrosis of animals in Group 1 (closed bars) and Group 2 (gray bars). The Chi-square test showed no statistically significant difference between the groups. **d**,**e** Representative examples of color coded, AHA 17-segment bullseye plots, showing the results of the analysis of regional wall thickness of the same animal (internal no. 12056) at T1 (**d**) and T2 (**e**). The asterisks indicate a segment that improved between T1 and T2. **f** Number of segments with unchanged, improved (+) and deteriorated (-) regional wall thickness of animals in Group 1 (closed bars) and Group 2 (gray bars). The Chi-square test showed a statistically significant difference between the groups.

**Supplementary Fig. 3.**
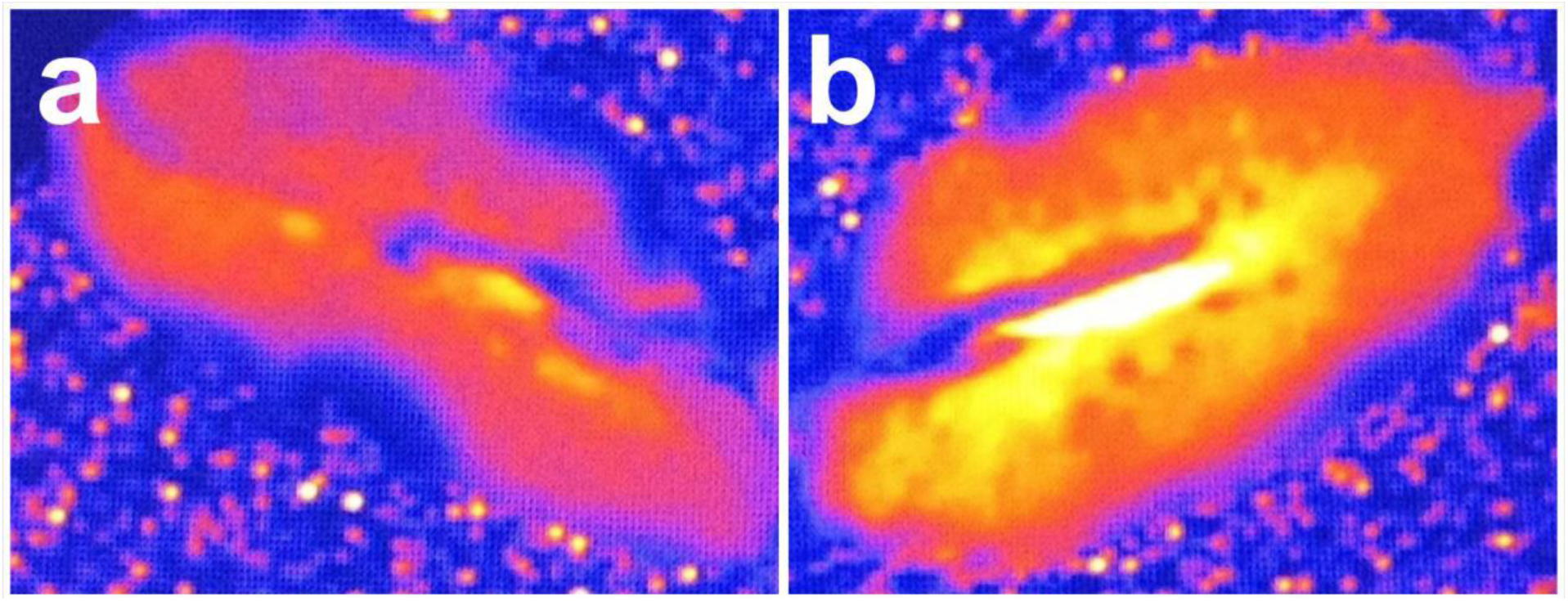
T2* magnetic resonance imaging^4,5^ of the right (**a**) and left (**b**) kidney of a pig with normal kidneys (Group 5) after delivery of autologous, adipose-derived stem cells (A-ASCs) labeled with iron-oxide particles into both renal arteries. The images are color-coded and represent R2* (reciprocal of T2*) in ms, ranging from blue (lowest; 0 ms) over yellow (middle) to red (highest; 100 ms). For the right kidney (**a**), during cell delivery the runoff of cells was blocked by an inflated balloon catheter. The delivery of cells into the left kidney (**b**) was performed without blockage. Due to the inflated balloon during cell delivery as in (**a**), cells became trapped between the inflated balloon and the capillary bed during initial delivery of the cell suspension. When cell delivery continued, an increased pressure resulted in the vascular compartment between balloon occlusion and capillary bed, enabling the cells to overcome the endothelial barrier and to extravasate into the parenchymal tissue (**a**), rather than being trapped only in the peripheral capillary bed (**b**). Increased iron retention in tissue resulted in shortening of T2*, which is reflected by the purple/dark red color, while low iron / cell content appears yellow. With the blocked injection in (**a**), a more homogenous distribution and larger number of cells in the kidney (indirectly indicated by the iron content as red color) was found compared to the non-blocked delivery in (**b**). Labeling of A-ASCs with iron-oxide particles was performed as previously described^6^. Briefly, 10-100 µl of 3×10^8^ encapsulated superparamagnetic microspheres (with diameters of 0.96, 1.63, 2.79 and 5.80 µm and consisting of a magnetite core coated with a divinyl benzene/styrene polymer; Bangs Laboratories, Fishers, IN, USA; or with a diameter of 4.5 µm and consisting of a magnetite core coated with a polystyrene polymer; Dynal Biotech, Oslo, Norway) were added to the growth medium, and incubated with the A-ASCs for 18 hr. COOH functionality was introduced to the surface of the particles, and a fluorescent dye was soaked into the polymer coating of some of the particles. To remove loosely bound and free particles, the cells were washed extensively. Then, the cells were released from the dish by incubation with trypsin. To remove additional free particles, the cells were pelleted and resuspended in growth medium at 1×10^6^/ml, and density centrifugation was performed with Ficoll-Paque PLUS (Amersham Biosciences, Piscataway, NJ, USA). To this end, 3 ml of Ficoll-Paque PLUS was added to 15-ml plastic Falcon tubes. Then, 3 ml of resuspended cells were layered on top of the Ficoll-Paque PLUS, with care taken to not disturb the interface. The tubes were centrifuged for 30 min at 400 g. After centrifugation, most of the free particles had pelleted to the bottom of the tube, and the cells remained at the interface of the Ficoll-Paque PLUS and the growth medium. The cells were removed with a glass pipette and washed with growth medium to remove the remaining Ficoll-Paque PLUS. For imaging purposes, the cells were transferred onto eight-well chamber slides (Lab-Tek II; NUNC, Rochester, NY, USA) (50 cells per well), and allowed to attach. Growth medium was made with 1 mM Gd-DTPA (Magnevist, Berlex) to shorten T1* in order to allow rapid MR imaging. Trypan blue exclusion tests were performed at each step to measure cell viability.

**Supplementary Fig. 4.**
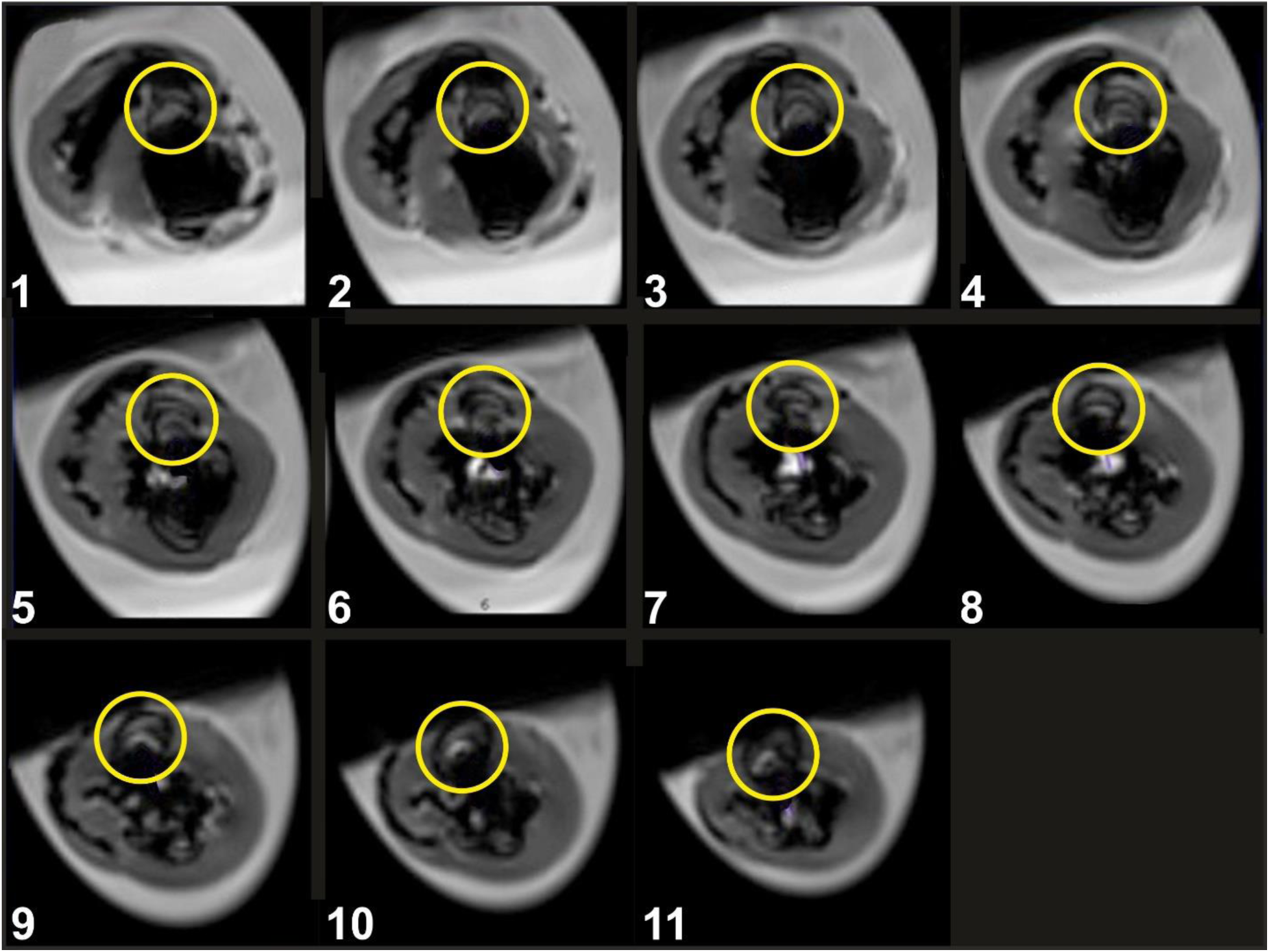
Short-axis T2* cardiac magnetic resonance (CMR) images showing the distribution of autologous adipose-derived stem cells labeled with iron-oxide particles after cell delivery through the left anterior descending (LAD) vein in a porcine heart without induction of myocardial infarction (Group 5 in the main text). Cells were labeled as explained in the legend of Suppl. Fig. 3 and were delivered in the same manner as described for animals in Groups 1 and 3 in the main text. Thirty minutes after cell delivery the animal was euthanized, and the heart was harvested and imaged with T2* CMR imaging. The signal distortion that was caused by the iron oxide particles taken up by the delivered cells is highlighted with yellow circles. The delivered cells were distributed in the anterior septal segments which corresponded to the site where myocardial infarction was experimentally induced by occluding the LAD artery of the animals in Groups 1-4 of the present study.

## SUPPLEMENTARY MOVIES

**Supplementary Movie 1** Cine sequence (obtained with steady-state free precession cardiac magnetic resonance (SSFP CMR) imaging and covering one full period of the cardiac cycle) of short axis, transversal images of the mid left ventricle of the heart of an animal in Group 1 (c.f. Table 1 in the main text), obtained at four weeks (time point T1) after experimental occlusion of the left anterior descending (LAD) artery by means of a balloon catheter for three hours at time point T0.

**Supplementary Movie 2** Cine sequence (obtained with steady-state free precession cardiac magnetic resonance (SSFP CMR) imaging and covering one full period of the cardiac cycle) of short axis, transversal images of the mid left ventricle of the heart of the same animal in Group 1 (c.f. Table 1 in the main text) as shown in Supplementary Movie 1, obtained at ten weeks (time point T2) after experimental occlusion of the left anterior descending (LAD) artery by means of a balloon catheter for three hours at time point T0, followed by the delivery of 18 × 10^6^ unmodified, autologous adipose-derived regenerative cells into the LAD vein (matching the initial LAD occlusion site) immediately after SSFP CMR imaging at T1.

**Supplementary Movie 3** Cine sequence (obtained with steady-state free precession cardiac magnetic resonance (SSFP CMR) imaging and covering one full period of the cardiac cycle) of short axis, transversal images of the mid left ventricle of the heart of an animal in Group 2 (c.f. Table 1 in the main text), obtained at four weeks (time point T1) after experimental occlusion of the left anterior descending (LAD) artery by means of a balloon catheter for three hours at time point T0.

**Supplementary Movie 4** Cine sequence (obtained with steady-state free precession cardiac magnetic resonance (SSFP CMR) imaging and covering one full period of the cardiac cycle) of short axis, transversal images of the mid left ventricle of the heart of the same animal in Group 2 (c.f. Table 1 in the main text) as shown in Supplementary Movie 3, obtained at ten weeks (time point T2) after experimental occlusion of the left anterior descending (LAD) artery by means of a balloon catheter for three hours at time point T0, followed by the delivery of saline (as control) into the LAD vein (matching the initial LAD occlusion site) immediately after SSFP CMR imaging at T1.

